# CRISPRi-mediated silencing of *racR* reveals molecular strategies for surviving lethal stress coupled to Rac prophage excision

**DOI:** 10.64898/2026.09.06.749655

**Authors:** Gargi Bindal, Devashish Rath

## Abstract

Cryptic prophages, abundant in bacterial genomes, often carry regulatory modules that rewire host physiology contextually. *E. coli* K-12 Rac prophage harbors a genetic module consisting of a RacR repressor that tightly represses the toxin genes ydaS/T. Dysregulation of *racR* and expression of YdaS/T lead to growth arrest, rapid loss of viability, and morphological defects. We investigated the physiological consequences of *racR* silencing and the host adaptive responses to lethal stress imposed by YdaS/T. We show that *racR* depletion establishes an inner membrane-centered stress characterized by selective hyperpolarization, increased permeability, and arrest of cytokinesis downstream of FtsZ-ring assembly, triggering copious filamentation. Transcriptomic analysis showed extensive metabolic reprogramming, revealing a coordinated, energy-conserving response marked by respiratory remodelling, repression of proton motive force-intensive pathways, and activation of global stress pathways. We show that recovery is accompanied by remarkable physiological heterogeneity with divergent filament fates, including lysis, persistent arrest, and recovery by loss of Rac. Thus, heterogeneous membrane dysfunction and metabolic states appeared to determine the balance between recovery and cell death. These findings support a model in which *racR* silencing activates a prophage-host stress circuit interlinking inner membrane energetics, activation of envelope stress, cell division control, and heterogeneous recovery.

## Introduction

Prophages, particularly cryptic prophages, which have lost one or more functions required for a productive phage cycle, such as excision, replication, or host lysis, create long-lived genetic reservoirs that can be inherited vertically, transferred horizontally, and reshaped by mutation, recombination, and selection over time. This process contributes substantially to bacterial genome plasticity, strain diversification, and ecological adaptation (1–3). Prophage genes can encode toxins, antitoxins, recombination systems, transcription factors, envelope proteins, metabolic functions, defense modules, and virulence determinants, allowing bacteria to acquire phenotypes that would be difficult to evolve through point mutations alone (4,5). In *Escherichia coli* K-12, cryptic prophages account for almost 4% of the genome, whereas in some pathogenic *Escherichia coli* (*E. coli)* strains, such as O157:H7 Sakai, the prophage burden can reach nearly 16% of the chromosome (6, 7). Prophages engage host transcriptional networks to enhance survival under osmotic, oxidative, and acidic stress, modulate biofilm formation, and influence antibiotic tolerance (8–10). A comprehensive understanding of how these genetic elements integrate with host cellular physiology and influence survival under adverse conditions, is essential. Furthermore, it is critical to investigate the consequences of dysregulation of phage genetic elements and the host cellular mechanisms such as metabolic reprogramming and activation of stress-adaptive survival pathways elicited in response.

The cryptic Rac prophage in *E. coli* K-12 encompasses approximately 23 kb of the genome and comprises 24 genes and five pseudogenes (11–13). Rac harbours genes involved in recombination, excision, toxin-antitoxin systems, and transcriptional control but does not produce infectious progeny. The Rac prophage has been linked to biofilm formation, stress adaptation, and bacterial fitness, depending on the genetic background and the environmental context (10). Previously, we established a toxin-repressor system in the Rac prophage involving a repressor, RacR, and two adjacent co-expressed toxins, YdaS and YdaT. Silencing of *racR* by CRISPRi activates the expression of *ydaS/T* genes, resulting in growth arrest, cell death, and drastic morphological changes (12). RacR binds multiple operator sites of the strong *P_ydaST* promoter, which drives the bicistronic *ydaS-ydaT* transcript and keeps it tightly repressed (14, 15).

Initial genetic studies with individual deletion of *ydaT* or *ydaS* under RacR-depleted conditions showed differential effects on filamentation and cell death, suggesting distinct functions of YdaS and YdaT (12). Recent studies have shown that YdaT accounts for most of acute toxicity and cell division disruption, whereas YdaS promotes Rac excision (11, 15–17). YdaT binds upstream of the host *rcsA* gene and activates its expression, thereby engaging the RcsA/RcsB envelope stress regulon. Consequently, YdaT expression reduces motility, promotes biofilm formation, alters sensitivity to membrane-disrupting conditions, changes susceptibility to certain phages, and causes filamentation in genetically susceptible host. While, imbalance of RacR or YdaT is reflected as rapid loss of viability and extensive morphological alterations, it is interesting to note that the cells eventually recover and resume normal growth as the lethal YdaT effect is counteracted by Rac prophage excision, presumably mediated by YdaS (16, 17). Defective replication of Rac ensures its loss in subsequent division cycles, providing an escape route from cell death.

While recent studies have shed light on the cascade of events following RacR depletion, many aspects require deeper investigation. For instance, the transient phenotype that is reversed after Rac excision has not been studied at the single-cell level. This is particularly important because Rac excision has already been shown to occur over generations, unlike the rapid timescale of lambda induction (16). Moreover, the processes contributing to heterogeneity in the *racR*-silenced population and concomitant YdaS/T expression remain to be thoroughly explored. Prior studies have shown that some cells die, some survive long enough to lose Rac, and recovered populations emerge after prolonged perturbation (16, 17). However, population recovery can occur through different mechanisms, including the reversal of filamentation in individual cells, selective outgrowth of less perturbed cells, genetic loss of Rac, or a combination of these processes. Therefore, it is pertinent to ask whether recovery reflects active physiological reversal or is only replacement by unaffected cells and to attempt to distinguish between these alternatives.

In this study, we performed physiological depletion of *racR* using the CRISPRi approach and mechanistically studied prophage excision and YdaS/T-driven stress on host cell physiology. We combined time-resolved measurements of Rac prophage excision with assays of membrane potential, membrane permeability, and live-cell morphology to define the events following the activation of the RacR regulatory module. This allowed us to probe YdaS/T-associated membrane and division phenotypes that unfolded during the same interval in which Rac was excised and lost. We specifically asked whether YdaS/T-induced filamentation was an irreversible state, whether membrane perturbation was an early and measurable component of the response, and how the bacterial population recovered from the initial physiological crisis. We demonstrated that YdaS/T toxicity is a dynamic cellular response rather than a simple terminal toxicity phenotype. We resolved the time course of Rac prophage excision in relation to host physiology and identified membrane hyperpolarization and inner membrane permeabilization as mechanistic explanations for the stress imposed by YdaT. We further show that individual filaments resume division and contribute to population recovery in real time. These results broaden our understanding of the ways in which the activation of a replication-deficient prophage can impose significant physiological stress and the host’s approach to mitigate these lethal effects.

## Materials and methods

### 1. Bacterial strains and plasmids

*E. coli* K-12 MG1655 served as the parental strain in experiments described in this study. The strains and plasmids used are listed in Tables S2 and S3 respectively. The oligonucleotides used are listed in Table S4. Bacterial strains were cultivated with aeration at 37°C in Luria-Bertani (LB) or M9-CA medium (1× M9 salts, 0.1 mM CaCl2, 2 mM MgSO4, 10 μg/ml thiamine hydrochloride, and 1% (w/v) casamino acids, supplemented with 0.4% glucose) depending on the experimental endpoint. When necessary, plasmids were maintained using kanamycin (50 μg/ml), ampicillin (100 μg/ml), streptomycin (50 μg/ml), tetracycline (25 μg/ml), and chloramphenicol (15 μg/ml). For FtsZ localization studies, the MG1655 strain was engineered to express an FtsZ-GFP fusion protein from the chromosomal locus, *lacI*. The *lacIqP208-ftsZ-gfp* was introduced into MG1655 by P1vir transduction, using BS001 (18) as the donor strain, as previously described (19). This strain was designated MG1655_FtsZ.

### 2. Expression of *ydaS/T* from native locus

The binding of the RacR to the native promoter of the *ydaS-ydaT* operon keeps the operon repressed. For YdaS/T protein expression, repression was relieved by knocking down *racR* expression using the Type I-E CRISPR-based CRISPRi established previously (12). Appropriate strains were transformed with pWUR400 (expressing Cascade from a T7 promoter), and pCRISPR plasmid expressing the P1 or O1 spacer containing crRNA from PLlacO-1 (IPTG-inducible) to introduce CRISPRi (12). Silencing was induced using 0.2% L-arabinose and 0.1mM IPTG at OD ∼0.1 as required. As IPTG causes overexpression of FtsZ and filamentation even in control cells, an IPTG-independent CRISPR-based silencing vector pCBG1.1 was used to induce *racR* silencing in the strain expressing FtsZ::GFP. To construct, pCBG1.1, Cascade operon was amplified from pWUR400 plasmid using Q5 High-Fidelity DNA polymerase (NEB) and assembled by Gibson cloning with two additional fragments: a custom-synthesized JW23119-crRNA cassette and the p15A ori fragment derived from the pdCas9 plasmid, which had been linearized by *BglII* and *XhoI* (20), there by generating the all-in-one CRISPRi vector, pCBG1.1. Silencing was induced with 1 µM aTc (Cascade) and 2.5 µM IPTG (FtsZ-GFP).

### 3. Nucleoid and membrane staining

To visualize nucleoids and cell membranes, cultures were harvested 5 h post-induction of *racR* silencing and co-stained with DAPI (1 µg/ml; excitation/emission, ∼358/461 nm) and FM4-64 (4 µg/ml; excitation/emission, ∼515/640 nm) for 10 min at room temperature in the dark. The stained cells were mounted on 1.5% agarose pads prepared in PBS and immediately imaged using a Leica THUNDER Imaging System with the appropriate filter sets. The images were processed using Fiji/ImageJ.

### 4. Flow cytometric analysis of cell morphology

Silenced and control cultures were diluted in PBS after 5 h induction, and FSC and SSC was recorded using a Partec CyFlow cytometer to assess changes in cell size and internal complexity associated with filamentation. Debris was excluded by appropriate FSC/SSC gating, and at least 100,000 events were collected per sample. The scatter plot was generated using FlowJo software.

### 5. FtsZ localization

To determine the consequences of YdaS/T expression on the critical step of cell division, MG1655_FtsZ strain was transformed with an all-in-one CRISPRi *racR*-silencing vector (pCBG1.1) and a non-target control plasmid. The cultures were grown overnight in minimal media and then sub-cultured into fresh minimal media supplemented with 1 μM aTc to induce CRISPRi and 2.5 μM IPTG to induce FtsZ-GFP expression. After five hours of induction, the cells were harvested, washed, fixed on an agarose pad, and visualized under a 100X magnification at an excitation wavelength of 488 nm and emission wavelength of 525 nm with a 490 nm dichroic mirror. Filaments exceeding 8 µm in length were selected for scoring distinct aspects of FtsZ::GFP distribution, such as the presence or absence of rings, the number of complete circumferential rings per filament, and the spacing between each ring, using appropriate plugins on the FIJI platform. The distribution of fluorescence intensity along the length of the filament was plotted against the fluorescence of the control cells in FIJI using the plot fluorescent profile feature (21).

### 6. Measurement of inner membrane potential

Inner membrane potential was assessed using the ratiometric probe 3,3′-diethyloxacarbocyanine iodide (DiOC₂(3)) with the BacLight Bacterial Membrane Potential Kit (Molecular probes), with some modifications (22,23). The optimized assay conditions included washing the cells in 1× PBS (pH 7.4) and treating them with 10 mM EDTA for 5 min to permeabilize the outer membrane. The cells were then washed twice and resuspended in an assay resuspension buffer containing 130 mM NaCl, 60 mM Na₂HPO₄, 60 mM NaH₂PO₄, 10 mM glucose, 5 mM KCl, and 0.5 mM MgCl₂ (pH 7.0). DiOC₂(3) was added to a final concentration of 30 μM, and samples were incubated in the dark at 37°C for 30 min, alongside an unstained control and control cells. A depolarized control was prepared by treatment with 5 μM CCCP for 15 min, followed by resuspension in assay buffer and staining with DiOC₂(3), as described above. The fluorescence was analyzed using a Partec CyFlow cytometer with 50000 events collected per sample. Green and red fluorescence were detected using 488 nm excitation with FL1 533/30 nm and 640 nm excitation with red FL4 670 nm long-pass, respectively. The FSC/SSC and gating parameters were set using the unstained control for each group and CCCP-treated depolarised control was used to set the voltage and gain. The FCS files were analyzed using FlowJo software and FCS express, and red and green mean fluorescence intensity (MFI), histograms, and dot plots were generated. All samples were examined using fluorescence microscopy to confirm the staining and polarization patterns.

### 7. Assessment of inner membrane permeability

Membrane permeability was assessed by monitoring ethidium bromide (EtBr) uptake and SYTO9/propidium iodide (PI) staining. For the EtBr uptake assay, silenced and control cultures were induced for 5 h, incubated with 6 μM EtBr, and fluorescence was recorded immediately and subsequently at 1-min intervals for 30 min using a Tecan multimode microplate reader (excitation, 545 nm; emission, 600 nm). The increased fluorescence intensity was interpreted as enhanced intracellular accumulation of EtBr, resulting from increased membrane permeability. Furthermore, permeability was evaluated by co-staining of 5 h induced silenced and control cells with SYTO9 (5 μM; excitation/emission, 485/498 nm) and propidium iodide (PI 1μg/ml; excitation/emission, 535/617 nm) according to standard protocols. Stained cells were mounted on 1.5% agarose pads prepared in PBS and immediately imaged at respective fluorescence channels sequentially to minimize spectral bleed-through. All images were processed using Fiji/ImageJ.

### 8. Assessment of outer membrane integrity

#### 8.1 Growth based assays

To assess susceptibility to the outer-membrane-impermeant antibiotic vancomycin, cultures were grown for 5 h in the absence of induction and then incubated at 37°C in the presence of sub-MIC concentration of vancomycin (0, 2, or 5 µg/ml) and inducers in 96 well plate; growth kinetics was monitored by measuring OD₆₀₀ at intervals of 30 min up to 30 h in a Tecan multimode reader with standard settings. To assess whether exogenous Mg²⁺ stabilizes the bacterial envelope and rescues the growth defect, cultures were induced for 5 h, serially diluted ten-fold, and spotted onto minimal medium agar plates containing the basal 2 mM MgSO₄ concentration or supplemented with MgSO₄ to final concentrations of 5 mM or 10 mM, along with the appropriate inducers. Plates were incubated under identical conditions at 37°C for 20 h before documenting bacterial growth.

#### 8.2 LPS quantification assay

Whole-cell lysates were prepared from *racR*-silenced and control cultures harvested 5 h post-induction to compare total lipopolysaccharide (LPS) levels. Briefly, cell pellets were resuspended in 10 mM HEPES (pH 7.4) containing 1 mM PMSF and 1× protease inhibitor cocktail, lysed by sonication, and clarified via centrifugation. Protein concentrations were determined using the Bradford assay, and equal amounts of total protein were used for LPS analysis. Samples were mixed with an equal volume of tricine sample buffer [100 mM Tris-HCl (pH 6.8), 24% (w/v) glycerol, 8% (w/v) SDS, 5% (v/v) β-mercaptoethanol, and 0.02% (w/v) bromophenol blue], boiled for 10 min, and digested with proteinase K (0.25 mg/ml) for 1 h at 60°C to remove proteins while preserving LPS. The digested samples were resolved on 12% tricine-SDS-PAGE gels and visualized by silver staining as previously described (24,25). LPS levels were quantified by densitometric analysis of lane intensity using Fiji/ImageJ.

#### 8.3 SEM analysis

Cultures were harvested 5 h post-induction, washed twice with PBS, and fixed with 2.5% glutaraldehyde for 1 h at 4 °C. Cells were washed and dehydrated through graded ethanol (10, 20, 30, 50, 70, 90, and 100%), followed by resuspension in 100% ethanol, mounting on stubs, air-drying, and gold coating. Samples were examined using a Zeiss scanning electron microscope operated at an accelerating voltage of 20 kV, a working distance of 7.0 mm, and a magnification of 30,000×.

### 9. Metabolic assays

#### 9.1 ATP Quantification

Intracellular ATP levels in *racR*-silenced and control cells were measured using the BacTiter-Glo Microbial Cell Viability Assay (Promega) according to the manufacturer’s instructions. This assay is based on the ability of firefly luciferase to catalyze the oxidation of d-luciferin in the presence of a magnesium salt and ATP, and the emitted light intensity is proportional to the ATP content, providing the status of cellular bioenergetics. For each measurement, 100 μL of bacterial cells was combined with 100 μL of BacTiter-Glo reagent in opaque walled 96 well plates, and the mixture was incubated at 37°C in the dark for 5 min. Luminescence was recorded on a TECAN Infinite 200 PRO multimode reader (1-s integration time) at 5-minute intervals over a 30 min time window to confirm signal stability. Extracellular ATP was quantified from cell-free supernatants obtained by centrifugation (16,000 × g, 2 min, 4°C) and filtration through a 0.22 μm filter to distinguish intracellular depletion from membrane leakage.

#### 9.2 Respiratory activity

Respiratory activity in individual cells of the YdaS/T-expressing population was assessed using the redox-sensitive fluorescent probe 5-cyano-2, 3-ditolyl tetrazolium chloride (CTC), which is reduced intracellularly by the electron transport chain in metabolically active cells to form insoluble, dark-red fluorescent formazan precipitates as described (26). Briefly, cells were harvested at the indicated time points post-induction, washed, and stained with freshly reconstituted 5 mM CTC for 1 h at 37°C in the dark. Following staining, the cells were washed, mounted on agarose pads, and imaged using a 100× objective (excitation 450 nm; emission 630 nm) on a Leica THUNDER imaging system.

### 10. RNA sequencing and differential gene expression analysis

Transcriptional profiling was performed in three groups: high silencing (crRNA P1), moderate silencing (crRNA O1), and control. Following a 5 h induction of CRISPRi, total RNA was extracted from three independent clones per condition, as described (27). RNA integrity and concentration were assessed, and ribosomal RNA was depleted using the Ribo-Zero rRNA removal system. Strand-specific sequencing libraries were prepared from rRNA-depleted RNA using the TruSeq Stranded Total RNA Library Prep Kit (Illumina) according to the manufacturer’s instructions and sequenced on an Illumina HiSeq 2500 platform in paired-end mode (2 × 150 bp). Paired-end reads were aligned to the *E. coli* K-12 MG1655 reference genome (GCF_000005845.2_ASM584v2) using HISAT2, and gene-level counts were generated using feature counts with reverse-stranded read counting to reflect strand-specific library chemistry. Differential gene expression analysis was performed using DESeq2 (v1.40.2) across three experimental groups: C1 (n = 3), P1 (n = 3), and O1 (n = 2). Dispersions were estimated using the parametric fit, differential expression was assessed using the Wald test, and P-values were adjusted for multiple testing using the Benjamini–Hochberg false discovery rate (FDR) procedure (28,29). For each pairwise comparison (O1 Vs C1, P1 Vs C1, and P1 Vs O1), differentially expressed genes (DEGs) were defined as those with an adjusted P-value < 0.05 and an absolute log2 fold change > 1, corresponding to a minimum two-fold change in expression. Functional enrichment analysis of the DEGs was performed using ShinyGO (v0.85.1), with separate analyses of the upregulated and downregulated gene sets from each pairwise comparison, against Gene Ontology (Biological Process, Cellular Component, Molecular Function) categories at an FDR threshold of 0.05 (30). Transcriptional regulatory interactions were obtained from the RegulonDB Network Regulator Gene dataset (31), and processed using custom Python scripts (Figure S12) to identify transcriptional regulators enriched in DEGs list belonging to each comparison group. The full parameters for enrichment, visualization, and regulatory network analyses are provided in Supplementary text section 1.

### 11. Quantitative reverse-transcription PCR

Total RNA was isolated using the RNASNAP method followed by DNase digestion. For subsequent quantitative reverse-transcription PCR (qRT-PCR) analysis, 1 µg of total RNA from each sample was used for cDNA synthesis (Themo: RevertAid First Strand cDNA Synthesis Kit) according to the manufacturer’s instructions. Quantitative PCR was performed using KAPA SYBR FAST qPCR Master Mix (2X) Kit, on a Roche light cycler using standard cycling conditions. Relative transcript abundance was calculated using the ΔΔCt method after normalization to 16S rRNA and *rpoS* genes (32). The Primer sequences are listed in Table S4.

### 12. Estimation of Rac prophage excision

To quantify the frequency of Rac prophage excision over the time course of *racR* silencing, cells were sampled at 3, 5, 8, and 16 h post-induction, along with a non-targeting control population expressing wild-type levels of YdaS/YdaT. At each time point, individual clones were isolated by plating on LB agar plates without inducers. From the resulting colonies, 35 clones from the *racR-*silenced population and 12 clones from the control population were selected for analysis. Genomic DNA was extracted from each clone, and qPCR (KAPA SYBR FAST qPCR Master Mix (2X) Kit) was performed using primers flanking the *ydaS/T* locus and *rpoS* as a single-copy reference gene (Table S4). The qPCR conditions were as follows: pre-denaturation step 95 ◦ C, 30 s and 40 cycles of 95 ◦ C for 10 s, annealing 55 ◦ C for 20 s and elongation at 72 ◦ C for 10 s, Tm 60 ◦ C to 95◦ C. The reaction was performed using a Roche Light Cycler 480. Each reaction was performed in technical triplicates and repeated independently at least twice. Melting curve analysis was performed to confirm the formation of specific products. For each clone, the presence or absence of the *ydaS/T* locus in the genome was determined from the absolute Ct value obtained in the sample that showed positive *rpoS* amplification under *racR*-silenced and control conditions. A positive melting curve profile was used to distinguish true amplification from spurious Ct values, which arose from very low template concentrations in the *racR*-silenced clones. This allowed the proportion of clones that had undergone Rac prophage excision to be calculated at each time point in both silenced and control populations.

### 13. Time lapse microscopy

For time-lapse microscopy of YdaS/T-expressing cells, overnight cultures were subcultured in minimal medium containing appropriate antibiotics and inducers to an OD_600_ of 0.1 to 0.2 (∼2 h) and loaded into a glass slide chamber. The slide chamber was prepared by fixing a Gene Frame (Thermo Cat#AB0577) on a standard glass slide. To mobilize the cells, 1.5% Low EEO agarose pads were prepared in minimal media supplemented with inducers. The agarose pad was cut into an H-shape to aerate the growing cells. Cells were loaded onto an agarose pad and maintained at 37°C in situ under the microscope objective throughout the imaging experiment. Images were captured every 5-10 minutes, for maximum of 5 h. Microscopy of cells grown on slides was performed using a Leica THUNDER imager equipped with temperature-controlled stage at 60× differential interference contrast (DIC) objective. Images and time-lapse videos were adjusted for brightness and contrast and processed into movie files using FIJI and cellSense software.

### 14. Statistical analysis

Statistical analysis were performed using GraphPad Prism 8.0.2. Quantitative data are presented as mean ± standard deviation (SD) from at least three independent biological replicates, unless otherwise stated. The median is presented for data for which normality could not be assumed. The specific statistical test used for each experiment, number of biological replicates (n), and exact P values or significance thresholds are indicated in the corresponding figure legends.

## Results

### 1. CRISPRi-mediated silencing of *rac*R induces pleiotropic effects on cellular physiology

The *racR* gene of the Rac prophage encodes a transcriptional repressor of the adjacent and divergently transcribed operon consisting of *ydaS* and *ydaT* genes. To dissect the physiological consequences of *ydaS/T* derepression from native promoters, we used a Type I-E Cascade – mediated CRISPRi platform to silence *racR,* which resulted in coordinated growth arrest within five hours, and was characterised by extensive cell filamentation, loss of viability, and cell lysis (12). We used three distinct crRNAs (P1, O1, and O2) to direct the Cascade complex to distinct positions within the *racR* coding sequence, positioned +3, +58, and +208 bp relative to the start codon, respectively (Figure 1 A), resulting in graded depletion of RacR levels. A scrambled non-targeting sequence in the control crRNA was used as a negative control. The effect of graded depletion of RacR on cellular morphology was studied using flow cytometry, in which forward scatter (FSC) served as an indicator of cell size. The FSC progressively increased in the O2, O1 and P1 populations relative to the non-targeting control, with a corresponding rightward and upward shift in the FSC/SSC density plots (Figure 1 B). This confirms that the different levels of filamentation achieved by each crRNA can be quantified using flow cytometry-based assay. This pattern was corroborated at single-cell resolution by directly measuring cell length from bright-field micrographs (Figure 1C). Control cells (n = 208) were short, whereas the three *racR*-silenced populations were heterogeneously elongated (O2, n = 213; O1, n = 316; P1, n = 173), with P1 showing the greatest median length and filaments up to ∼100 µm (Figure 1C). The cells reached approximately 10–20 times their normal length without division, with no significant change in width.

**Figure 1.**
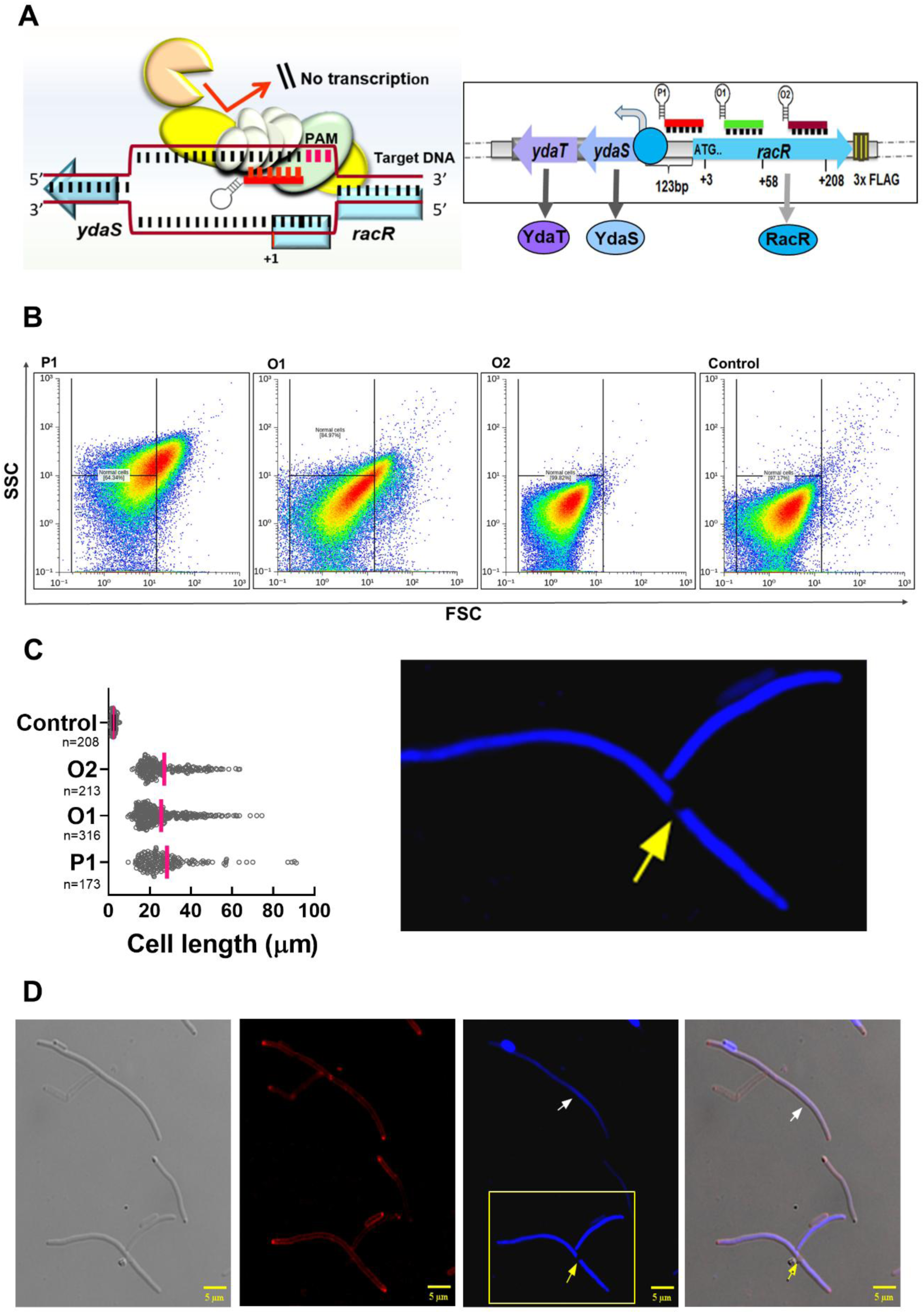
CRISPRi targeting of *racR* using a type I-E Cascade complex induces pleiotropic changes in *E. coli* cell physiology. Also see Figure S1. (A) Schematic of the CRISPRi system and target sites in *racR* locus. Left: Schematic of the type I-E Cascade complex binding target DNA to mediate transcriptional interference. Right: Linear map of the *racR* coding sequence showing three crRNA target sites (P1, O1, O2) located at +3, +58 and +208 bp relative to the start codon (B) FSC versus SSC density plots for mid-exponential phase *E. coli* populations expressing a non -targeting control crRNA or *racR* targeting crRNAs (P1, O1, O2). *racR* silencing produced a crRNA-dependent, progressive increase in FSC (O2 < O1 < P1), consistent with cell filamentation; representative density plots are shown; n = 3 biological replicates per condition. (C) Quantification of cell length from bright field micrographs. Density plots of measured cell lengths for control (n = 208), P1 (n = 173), O1 (n = 316) and O2 (n = 213), images pooled from three independent biological replicates. Red lines indicate median cell length. *racR* silenced populations showed heterogeneous elongation; and the cell length distributions differed significantly from the control for each crRNA (Kruskal–Walli’s test with Dunn’s multiple-comparison correction, p < 0.001 for each comparison). (D) Representative fluorescence micrographs of *racR* silenced cells stained with DAPI (blue, DNA) and FM4-64 (red, membrane). Elongated filaments with continuous nucleoid staining along the cell length (white arrowheads), and filaments exhibiting discrete DNA-free gaps interspersed between nucleoids (yellow arrowheads, zoomed to show gaps), indicating irregular nucleoid segregation or chromosomal partitioning defects. Membrane staining confirms an uninterrupted plasma membrane along the filaments. Micrographs are representative of three independent experiments.

To further investigate the impact of YdaS/T expression on cellular physiology, *racR*-silenced cells were stained with DAPI (DNA) and FM4-64 (membrane) dyes (Figure 1D; Figure S1). This revealed a spectrum of subpopulations, including filaments with continuous nucleoids and others exhibiting discrete gaps, indicative of irregular nucleoid segregation. These observations suggest that chromosome replication is uncoupled from septation in a subset of the cell population. A fraction of the cells retained normal rod morphology interspersed among the filaments, indicating that *racR* silencing did not drive the entire population into filamentation with equal kinetics or severity. Membrane staining with FM4-64 confirmed that the plasma membrane remained continuous and intact along the entire length of the elongated cell. However, the heterogeneity evident within the *racR*-silenced population raises the question of whether these cells represent a single uniformly arrested state or several mechanistically distinct subpopulations. To resolve this and determine whether the observed growth arrest constitutes an irreversible endpoint or a reversibly arrested state, we characterised FtsZ-GFP ring formation, membrane integrity and permeability, and viability in the *racR*-silenced population, mostly at single-cell resolution.

### 2. FtsZ ring assembly is disrupted but not abolished in *racR* silenced cells

To determine whether filamentation elicited by *racR* silencing reflects a failure of cell division and, if so, at which stage that failure occurs, we examined the localization of the essential division protein FtsZ using a chromosomally integrated *ftsZ-gfp* translational fusion at the *lacI* locus, which was induced by 2.5µM IPTG. The original two-plasmid CRISPRi platform required 0.1 mM IPTG to induce the expression of crRNA (12), and as both the fluorescent FtsZ fusion and the crRNA targeting *racR* were induced by IPTG, the higher IPTG concentration (0.1 mM) needed to induce crRNA expression also drove FtsZ-GFP overproduction, the non-targeting control cells became filamentous, irrespective of RacR depletion (33). To decouple the two induction regimes, we placed the crRNA array under the constitutive J23119 promoter and combined it with an aTc-inducible Cascade module to yield a single silencing vector, pCBG1.1 (Figure S2). *racR*-silenced filaments showed a heterogenous FtsZ-GFP distribution, including, multiple rings, dispersed foci, and diffused fluorescence (Figure 2A). Normal rod cells showed discrete Z-rings symmetrically positioned at the cell midpoint (inset in Figure 2A) consistent with canonical Z-ring assembly and Min system-mediated exclusion of the ring from the cell poles (34). Notably, a small subset of filaments exhibited constriction sites that appeared capable of progressing to successful cytokinesis, ultimately giving rise to normal rod-shaped daughter cells upon completion of division (Figure 2B). Among the filaments containing FtsZ rings, the mean number of rings per filament was approximately 2.14 and increased with filament length (Figure 2C), indicating that extended filaments frequently marked multiple putative division sites but failed to complete constriction at most of them. Quantitative analysis of 1,341 fluorescent filaments revealed that 785 filaments (59%) harboured at least one distinct circumferentially complete FtsZ ring, whereas the remaining 556 filaments (41%) displayed diffuse fluorescence without any organized ring structure (Figure 2D). The spatial arrangement of these rings was variable. Some filaments exhibited approximately equidistant ring spacing, consistent with Min-driven recovery from cell cycle arrest in filamentous *E. coli* (35), whereas others showed uneven positioning or clustering (Figure 2E). This result indicates that cell division could not progress further, either due to alterations in the inner membrane characteristics or inhibition of peptidoglycan synthesis and other cytokinesis steps. The filaments that showed no discrete rings were not due to the depletion of cellular FtsZ, as diffuse fluorescence was observed. Accordingly, the 41% of filaments without rings most plausibly represented a population comprised of a mix of cells with different possibilities, such as permanently growth-arrested cells, which would eventually die, or cells already dead, or in which ring nucleation was not achieved for reasons not known yet, or in which assembled rings disassembled in the presence of YdaS/T proteins.

**Figure 2.**
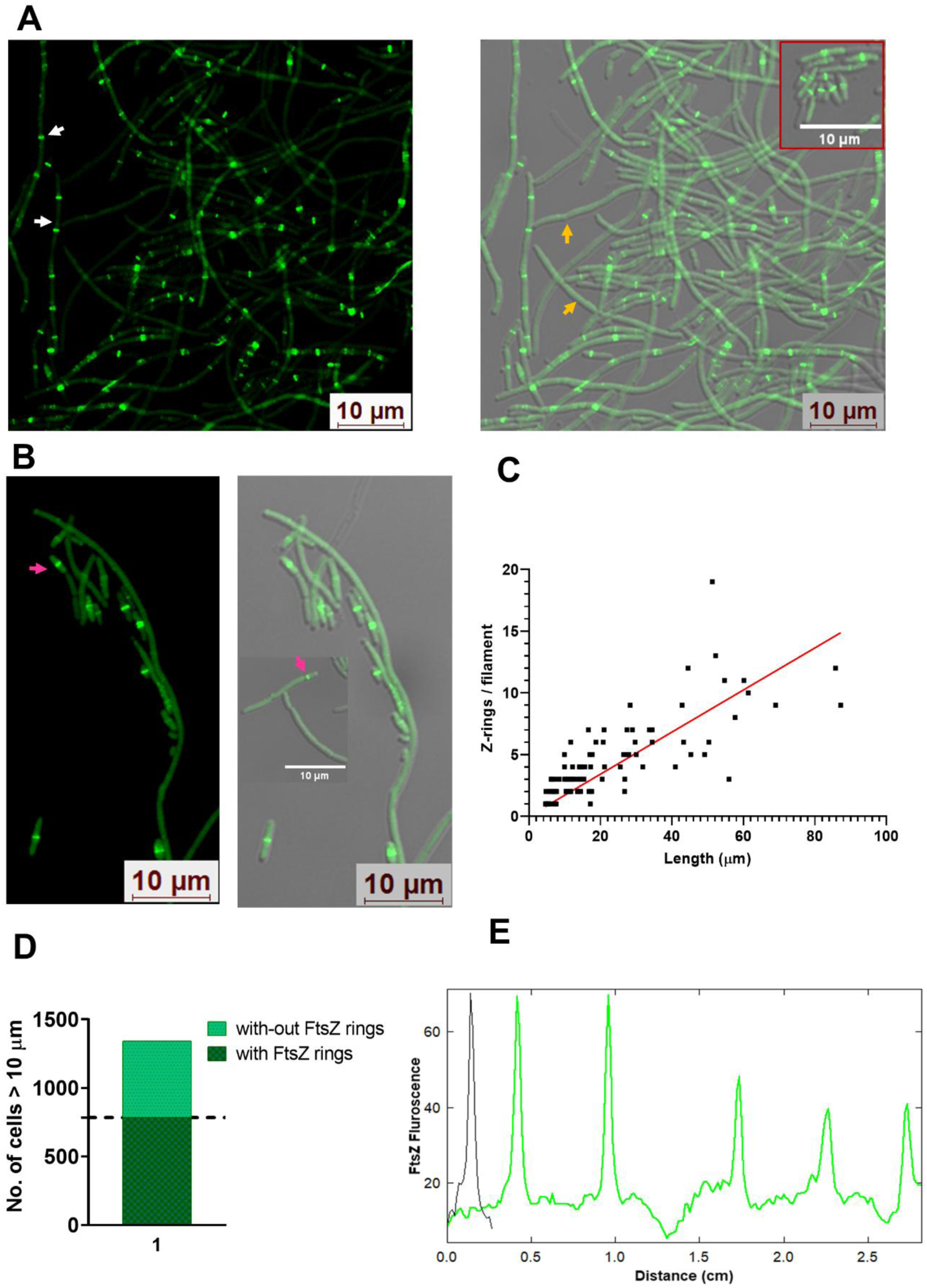
*racR* silencing uncouples FtsZ-ring assembly from cytokinetic completion. Also see Figure S2. *racR* was silenced using an all-in-one CRISPRi vector; knockdown was induced with 1 μM aTc, for 5 h prior to imaging. (A) Representative micrographs of FtsZ–GFP localization in *racR* silenced cells. Normal rods displayed a single, circumferentially complete Z-ring positioned at mid-cell (red box). The filaments showed heterogeneous FtsZ distribution, including multiple discrete circumferential rings along the length of individual filaments (white arrowheads) and filaments with diffuse, non-organized cytoplasmic FtsZ–GFP signal and no discernible ring (yellow arrowhead). (B) Representative *racR* silenced filaments showing FtsZ rings at putative division sites (pink arrowheads), indicating sites of eventual constriction and resolution into rod-shaped daughter cells. (C) Scatter plot of filament length (μm) versus number of FtsZ rings per filament (n = 104 filaments) with linear regression fit (red line) shown for visualization. Black squares indicated the number of FtsZ-GFP rings in filaments at the time of imaging. Filament length was significantly and positively correlated with ring number (Spearman’s rank correlation, rₛ = 0.81, P < 0.0001, two-tailed), indicating that longer filaments accumulated proportionally more FtsZ rings. (D) Quantification of FtsZ ring occurrence across individual fluorescent filaments (n = 1,341 filaments pooled from three independent biological replicates). 785 filaments (58.6%) contained at least one circumferentially complete FtsZ ring, whereas 556 filaments (41.4%) exhibited diffuse FtsZ–GFP fluorescence with no organized ring structure. Data are presented as total number of total filaments scored across the three biological replicates. (E) Line profiles of FtsZ:GFP fluorescence along representative filaments (green) compared with a normal rod-shaped cell (black), showing multiple fluorescence maxima in filaments versus a single division-site peak in rod-shaped cells.

These observations suggest that *ydaS/ydaT* activation did not arrest growth in an undefined manner but produced a specific division-competent-yet-division-incompetent state in which cells retained assembled FtsZ rings while failing to complete cytokinesis. This observed pattern of FtsZ distribution is qualitatively distinct from other known cell division inhibitors, such as the e14 prophage inhibitor YmfM/SfiC, which acts by direct sequestration of the FtsZ protein (36) or a dedicated anti-polymerization mechanism analogous to that of SulA (37). The presence of multiple mislocalized rings indicates a breakdown in the coordination between division site selection and downstream processes such as membrane constriction and cell wall synthesis. We interpret this as an abortive state of cell division, which we hypothesize is a consequence of the perturbed inner membrane physiology that accompanies *ydaS/T* derepression rather than a direct action on FtsZ. As the literature suggests that the characteristics of the inner membrane and energy availability are critical for proper divisome function, septal constriction, and anchoring of membrane-associated proteins in *E. coli* (38, 39), we next investigated these aspects.

### 3. YdaS/YdaT expression is associated with inner membrane hyperpolarization and increased membrane permeability

The proton-motive force (PMF) of the bacterial inner membrane, comprising an electrical component (ΔΨ, interior negative) and a chemical component (ΔpH), governs ATP synthesis, active transport, flagellar rotation, and critical cellular processes, such as anchoring of transport proteins, recruitment of cell division apparatus, PG synthesis, and cytokinesis (40,41). Because a perturbation of ΔΨ would be expected to propagate to each of these processes, the electrical state of the inner membrane was first examined as a direct biophysical readout of YdaS/T activity. The membrane potential was quantified by flow cytometry using the voltage-sensitive dye DiOC_2_ (3), which accumulates in cells in a ΔΨ-dependent manner. The dye is concentration-dependent and shifts from green to red in cellular compartments; thus, a red-to-green mean fluorescence ratio > 1 indicates increased inner membrane potential, independent of cell size (22). The protonophore CCCP, which collapses the PMF by transporting protons (H⁺) across the membrane, was included as a depolarisation control, and control cells provided the baseline membrane potential of the population at different stages of induction.

The expression of YdaS/T through *racR* silencing resulted in inner membrane hyperpolarization at 5 h post-induction. Quadrant analysis of the DiOC₂(3) FL1/FL2 dot plot showed that more than 60% of silenced cells exhibited elevated red fluorescence, suggesting that hyperpolarization was widespread across the population (Figure 3A; left panel). Representative micrographs of silenced cells displayed visible accumulation of DiOC_2_(3) red fluorescence compared to control cells (Figure 3A; right panel). The mean red/green ratio increased to 1.8 ± 0.3 in *racR*-silenced cells (Figure S3 A; additional micrographs shown in Figure S3 B). Flow cytometric analysis of the same cultures showed a corresponding rightward shift in the red/green fluorescence ratio distribution of silenced cells relative to control in representative overlaid histograms (Figure 3B). The proportion of cells within the high-ratio (M1) gate was significantly greater in *racR*-silenced cultures than in control. Together, these observations indicated that *racR* silencing increases the proportion of cells with elevated DiOC_2_(3) red/green ratios; critically, pre-treatment with the protonophore CCCP abolished the signal, demonstrating that the increased ratio reflects membrane hyperpolarization rather than altered dye uptake or cell-size artifacts.

**Figure 3.**
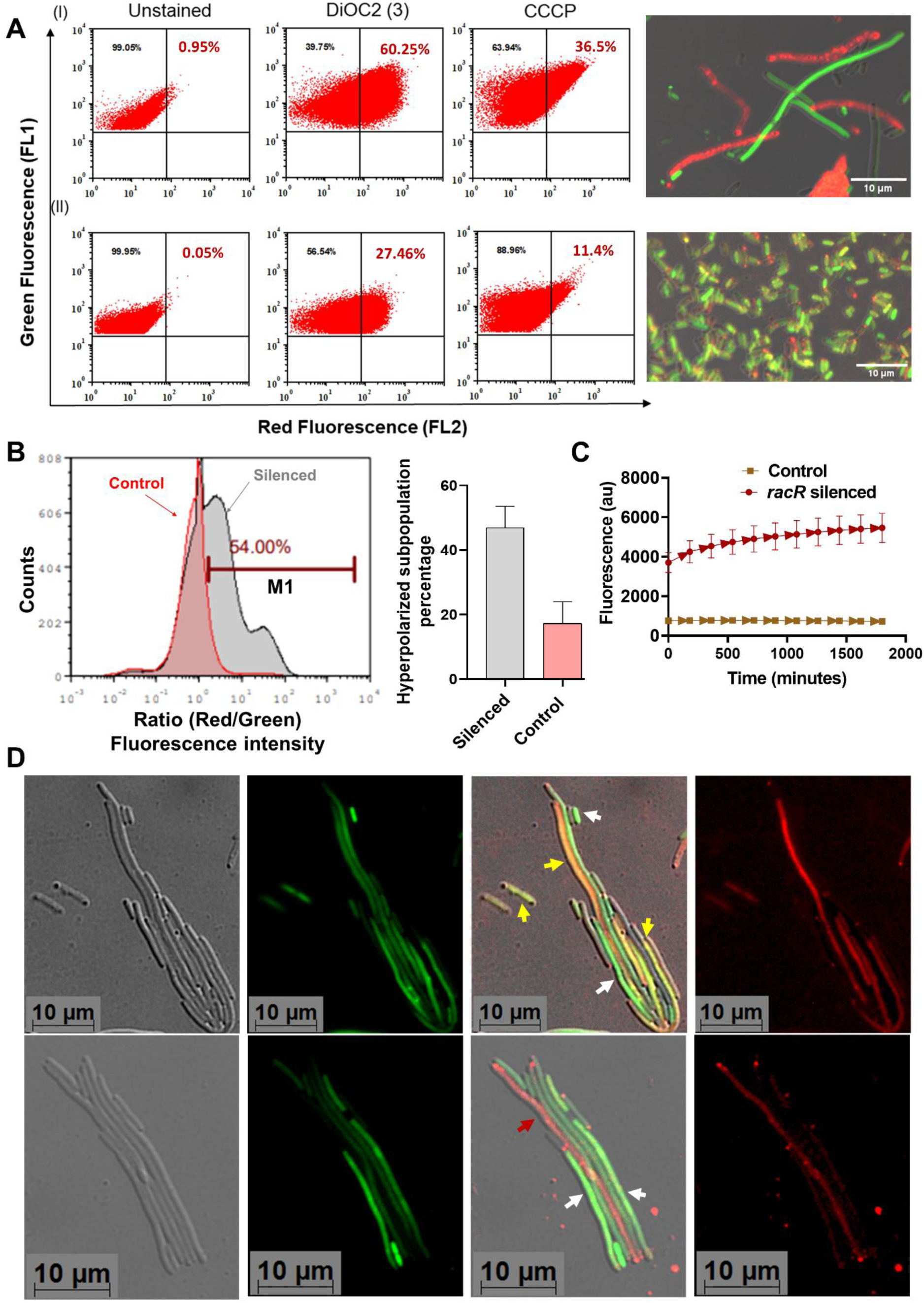
*racR* silencing hyperpolarizes the inner membrane and increases permeability in a subset of filaments. Also see Figure S3. (A) Flow cytometery-based measurement of membrane potential using DiOC_2_(3) staining 5 h after *racR* silencing. Representative dot plots of green fluorescence (FL1, cell size indicator) versus red fluorescence (FL2, membrane potential indicator) for (I) *racR*-silenced cells and (II) non-targeting control cells. Left: Unstained cells, middle: stained with DiOC_2_(3), right: treated with CCCP prior to DiOC_2_(3) staining as a control for assay sensitivity. Quadrant percentages indicate the proportion of cells hyperpolarized, with FL2 population reflecting increased red fluorescence characteristic of membrane hyperpolarization (n=3). Representative merged micrographs of *racR*-silenced cells and controls (scale bar, 10µm). (B) Representative overlaid flow cytometry histograms of the red/green fluorescence ratio for control (red) and *racR*-silenced (grey) populations, with the M1 gate boundary indicated. The bar graph shows the percentage of cells within the high-ratio (M1) gate. (Plotted as mean ± SEM; n = 3; unpaired two-tailed Student’s t-test, p <0.011). (C) The inner membrane is permeable to Ethidium bromide (EtBr). The *racR* silenced population rapidly accumulated Etbr compared to the control (data are mean + SEM, n=6, two-way ANOVA, p <0.001). (D) Dual staining with SYTO9 (green, permeant to all cells) and PI (red, excluded by cells with intact membranes) to distinguish viable from membrane-compromised filaments within the racR-silenced population. Representative micrographs showing SYTO9⁺/PI⁻ filaments with preserved membrane integrity (white arrowheads), SYTO9⁺/PI⁺ filaments with compromised membrane (yellow arrowheads), and SYTO9⁻/PI⁺ cells consistent with loss of membrane integrity (red arrowheads). Images are representative of three independent biological replicates.

Depolarization is the expected signature of pore-forming or ionophoric toxins, including the inner membrane toxins HokB and TisB, which dissipate ΔΨ by forming ion-conducting channels (42, 43). Hyperpolarization is the opposite polarity, which is contributed by an impairment of proton re-entry, for example, a restriction of flux through the F_1_F_0_-ATP synthase, while the respiratory chain continues to pump protons outwards, so that cell becomes more negative from inside and accumulates potential above its physiological levels. Furthermore, to establish whether hyperpolarization is accompanied by a loss of inner and outer membrane permeability, a panel of complementary permeability assays was employed in the silenced population. Inner membrane integrity was first probed with ethidium bromide (EtBr), a cationic DNA-intercalating dye whose entry is restricted by an intact, energized inner membrane and active efflux from the cell. At 5 h post-induction, *racR*-silenced cells accumulated significantly more EtBr than non-targeting control (Figure 3C), indicating increased inner membrane permeability. The increased EtBr signal could be due to a true barrier defect or efflux inhibition. This was further investigated by staining with SYTO9/PI dyes. Propidium iodide (PI) stains only permeable cells and is excluded by cells with an intact inner membrane, whereas SYTO9 stains only cells with an intact inner membrane. PI uptake was observed in approximately 58% of filaments (Figure 3D). Differential SYTO9/PI staining resolved this heterogeneity at the single-cell level, with PI-positive filaments (with a compromised inner membrane) coexisting with SYTO9-only filaments that retained an intact barrier and viability. This membrane integrity heterogeneity mirrored the observation of FtsZ rings in a similar percentage of filamentous cells, and together, the two readouts support the hypothesis that a population comprises both irreversibly compromised and physiologically recoverable filaments rather than a uniform terminal state.

In contrast to the inner membrane, the outer membrane remained largely intact, as confirmed by multiple assays. First, supplementation of the medium with Mg^2+^ (up to 10 mM), which stabilizes the outer membrane by bridging adjacent lipopolysaccharide (LPS) molecules and rescues outer membrane defects (44), did not alter growth or rescue the phenotype across the concentrations (2mM, 5mM and 10mM) tested (Figure 4A), indicating that the outer membrane was not compromised structurally. Consistent with this, LPS isolated from *racR*-silenced and control cells was present at comparable levels (Figure 4B), indicating that total LPS abundance remained unaltered despite perturbation of the inner membrane. The surfaces of the silenced cells were examined by SEM, which revealed smoother and poorly defined (fuzzy) cell boundaries in *racR*-silenced cells compared to the sharply defined and well-contoured surfaces of the control cells, suggesting alterations in cell envelope architecture (Figure 4C). However, sensitivity to vancomycin, whose entry into gram-negative cells depends on a permeabilized outer membrane, was similar in control and *racR*-silenced cells indicating that the outer membrane permeability was not substantially altered (Figure 4D). Taken together, these observations demonstrate that YdaS/T selectively compromises the inner membrane while leaving the outer membrane and its LPS architecture largely intact albeit with small changes in the cell envelope, a pattern that distinguishes this toxicity from generalized envelope stress responses that perturb both membranes simultaneously (45).

**Figure 4.**
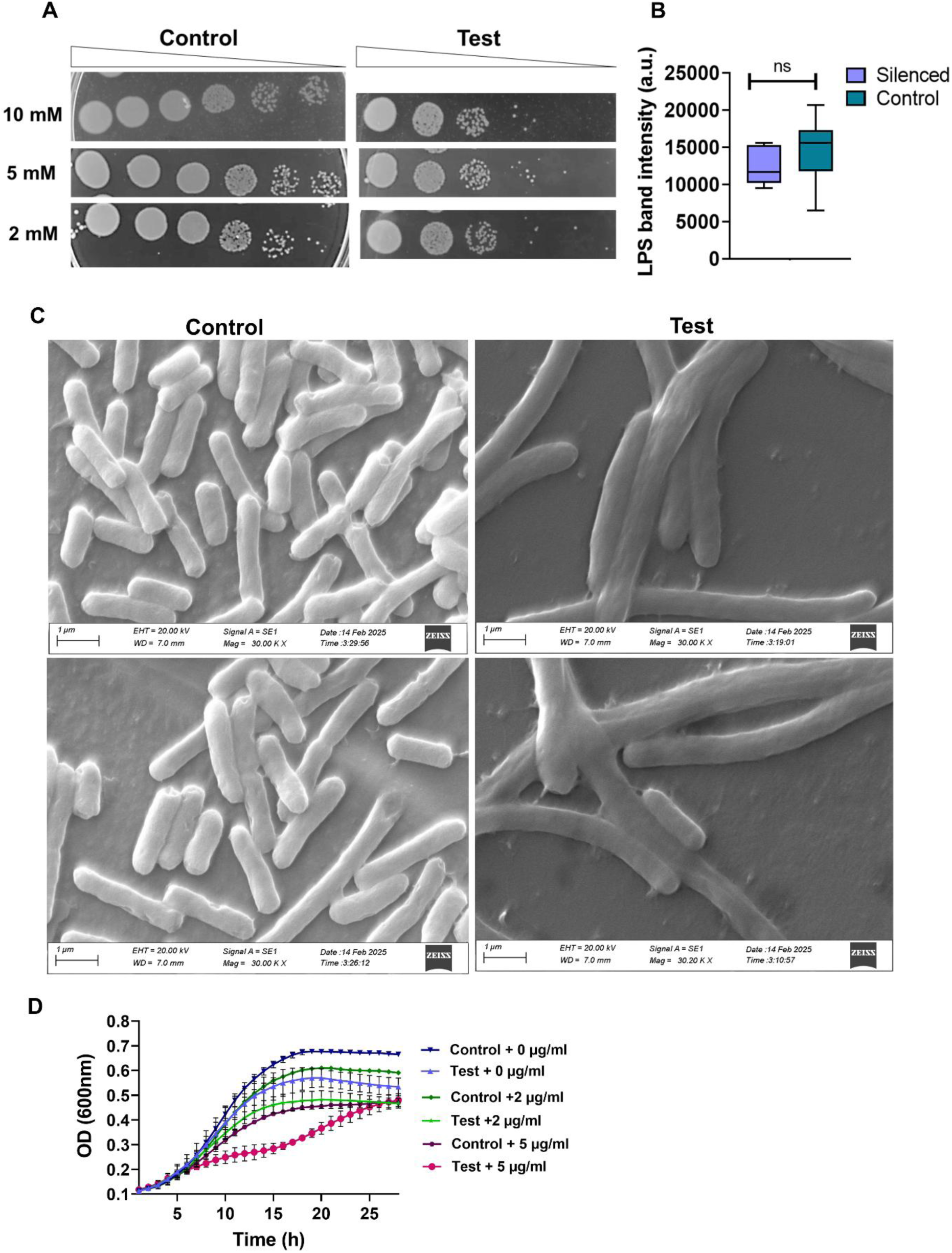
Outer membrane is unaffected in *racR* silenced population. (A) Spot assay on minimal media plates supplemented with 2mM, 5mM and 10 mM Mg+2 did not show improved growth survival of silenced cells as compared to the control (B) Bulk lipopolysaccharide content is not significantly altered by *racR* silencing. Total LPS was quantified by densitometric analysis of silver-stained tricine-SDS-PAGE gels. Box-and-whisker plots showing LPS band density for *racR*-silenced samples and controls (*n* = 9); horizontal line indicates the median; boxes span the interquartile range (Mann–Whitney *U* test, p <). (C) Representative SEM images showing fuzzy, smooth boundaries in *racR*-silenced cells. (D) Growth assay to determine sensitivity to Vancomycin. *RacR*-silenced cells (test) and control cells were exposed to indicated concentrations of vancomycin and growth was monitored for 30 h. Data represent mean ± SD from n = 3 independent biological replicates.

Collectively, these findings demonstrate that YdaS/T selectively perturbs the inner membrane, resulting in membrane hyperpolarization and increased inner membrane permeability while preserving the integrity of outer membrane. Given the pivotal role of the cytoplasmic membrane in respiration and ATP synthesis, we investigated the functional consequences of these perturbations on cellular bioenergetics and electron transport chain. Moreover, as a substantial proportion of the filamentous cells remained PI-negative, we next sought to determine whether these apparently membrane-intact filaments were metabolically active or inactive.

### 4. Filaments maintain active respiration despite ATP depletion

To assess the impact of membrane perturbation on cellular metabolism, we employed both population-level measurements of cellular ATP and 5-cyano-2, 3-ditolyl tetrazolium chloride (CTC) dye-based method to evaluate bacterial respiratory activity at the single-cell level (46). YdaS/T expression was induced for five hours, and ATP level was measured. For the first assay, the cells were treated with BacTiter-Glo reagent, which lyses the cells and catalyzes the substrate in the presence of ATP, producing a luminescent signal proportional to the cellular ATP content (Figure 5A). Compared to the control, the induced cells exhibited significantly reduced ATP levels, accompanied by elevated ATP levels in the filtered supernatant. This is consistent with our observation of inner membrane permeabilization, suggesting that disruption of membrane integrity contributes to ATP efflux and transient energy depletion in the internal milieu of bacterial populations. While population-level ATP measurements suggest impaired cellular energetics, they provide limited insight into the metabolic heterogeneity within the induced population.

**Figure 5.**
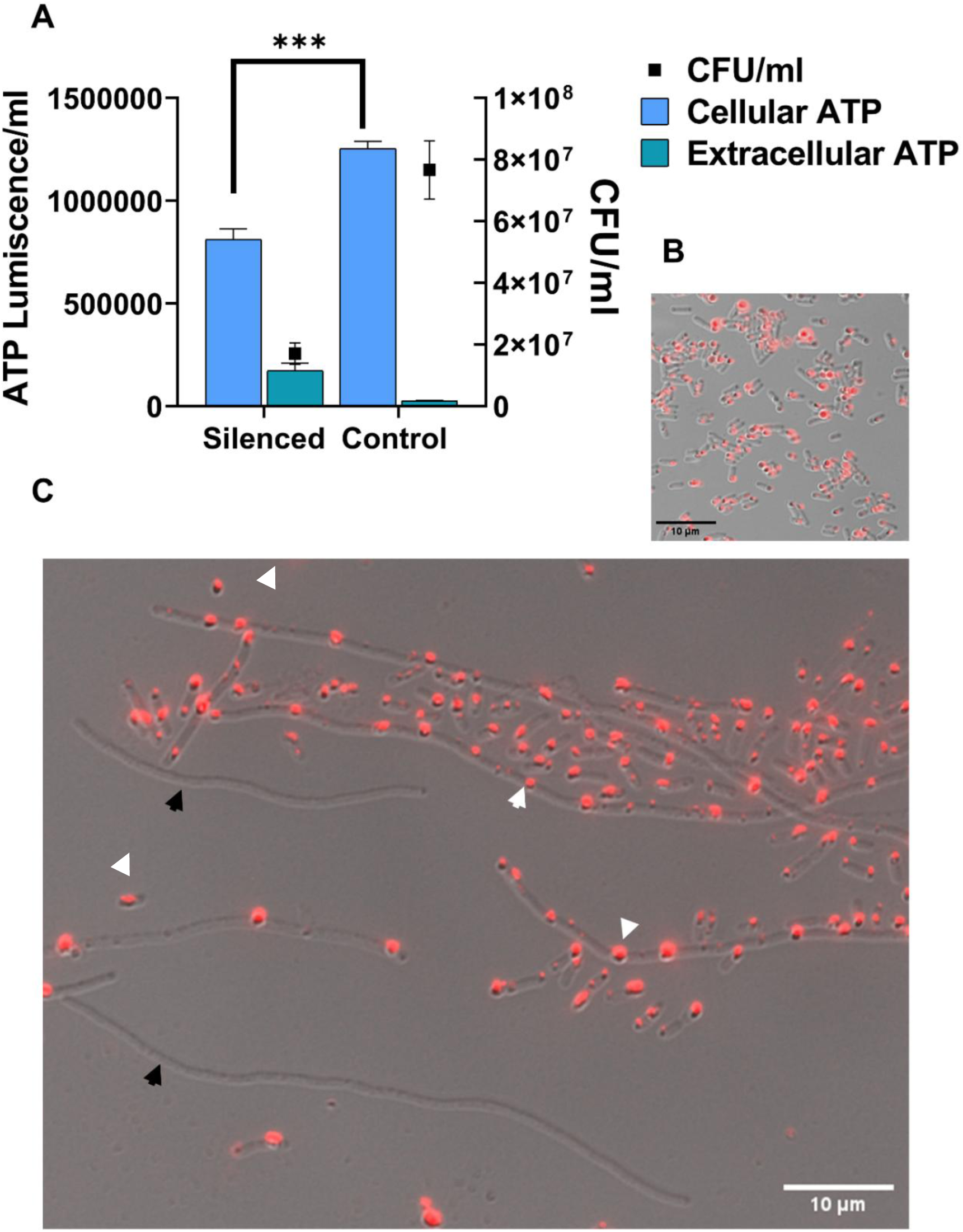
YdaS/T expression disrupts cellular ATP homeostasis and generates metabolically heterogeneous filamentous cells. Also see Figure S4. (A) ATP levels (bars, left y-axis) and viable counts (CFU/mL; squares, right y-axis, log₁₀ scale) were measured in control and *racR*-silenced cells. Intracellular ATP levels were significantly decreased, whereas extracellular ATP levels in the cell free supernatant medium increased, indicating membrane dysfunction and ATP leakage in silenced population. Data are presented as mean ± SD from n = 3 independent biological replicates. Statistical significance was determined using an unpaired two-tailed Welch’s t-test (***P < 0.001). (B) Representative images showing CTC (5-cyano-2,3-ditolyl tetrazolium chloride) reduction in normal cells. Scale bar, 10 µm. (C) Representative images showing CTC staining of silenced population showing heterogeneous respiratory activity. White arrows indicate filaments containing CTC formazan crystals (respiratory active), whereas black arrows indicate filaments with no detectable CTC reduction (respiratory inactivity). Scale bar, 10 µm.

Next, we employed the redox-sensitive CTC dye, which is reduced by cellular electron transport activity to form an insoluble, red fluorescent formazan precipitate that can be detected by fluorescence microscopy. This allows readout of respiratory activity of individual cells in heterogeneous populations. The control cells exhibited uniform active respiration throughout the population (Figure 5B). The RacR depleted population revealed substantial heterogeneity in respiratory activity within filamentous cells (Figure 5C; Figure S4). Filamented cells exhibited varying levels of red fluorescence; some demonstrated strong fluorescence, suggesting active electron transport and respiration, while others exhibited no detectable signal. This heterogeneity aligns with the observed variability in membrane integrity and suggests that *ydaS/T* expression induces a spectrum of metabolic states rather than a uniform response.

### 5. Global transcriptional remodelling defines the stress adaptive response to YdaS/T expression

To determine how the transcriptional changes underlying the observed filamentation, membrane defects, and cellular heterogeneity were coordinated at the global level in the *racR* silenced population, RNA sequencing was performed at two different silencing depths, designated P1 (high) and O1 (moderate), along with a control (C) without silencing. Principal component analysis of the expression data separated the three conditions, while biological replicates were clustered together, ensuring a high concordance (Figure S5). A roughly equal number of annotated genes was found to be expressed in each sample (FPKM>0). In total, 765, 418, and 662 DE genes were identified in the P1/C, O1/C, and P1/O1 comparison groups, respectively (Figure 6: upper left panel; Supplementary data I for the complete list of DE genes across different groups). Analysis of genes that were uniquely or commonly altered across the groups and the distribution of fold changes revealed widespread genome-wide transcriptional remodelling (Figure S6). Many genes were either induced or repressed in a silencing-dependent manner, thereby confirming progressive regulatory reprogramming under high and moderate silencing conditions. Common genes shared across conditions are shown in the heat map (Figure S7), and selected DE genes were validated by RT-qPCR (Figure S8). *racR* transcript abundance that served as an internal control was reduced by approximately 20-fold and 11-fold under P1 and O1 conditions, respectively. A strong derepression of genes present in the Rac prophage was observed (Figure 6: lower right panel). The adjacent toxin genes *ydaT* and *ydaS* were among the strongly induced transcripts genome wide, increasing by more than 100-fold in both conditions. Neighbouring Rac genes, including *racC, recE, recT, ralR, xisR, intR, kilR, ydaE, ydaG, and ydaU–W*, exhibited the same expression pattern (Table S5: A). Pathway enrichment analysis was performed to characterize the secondary physiological responses induced by *racR* silencing. The enriched pathways for each comparison are detailed in Supplementary text and presented in Figures S9–S11 (Supplementary Data II). Furthermore, regulon enrichment analysis based on RegulonDB annotations was performed to identify groups of DE genes under common transcriptional regulation (Figure S12; Supplementary Text, Table S5:E ; Supplementary Data II).

**Figure 6.**
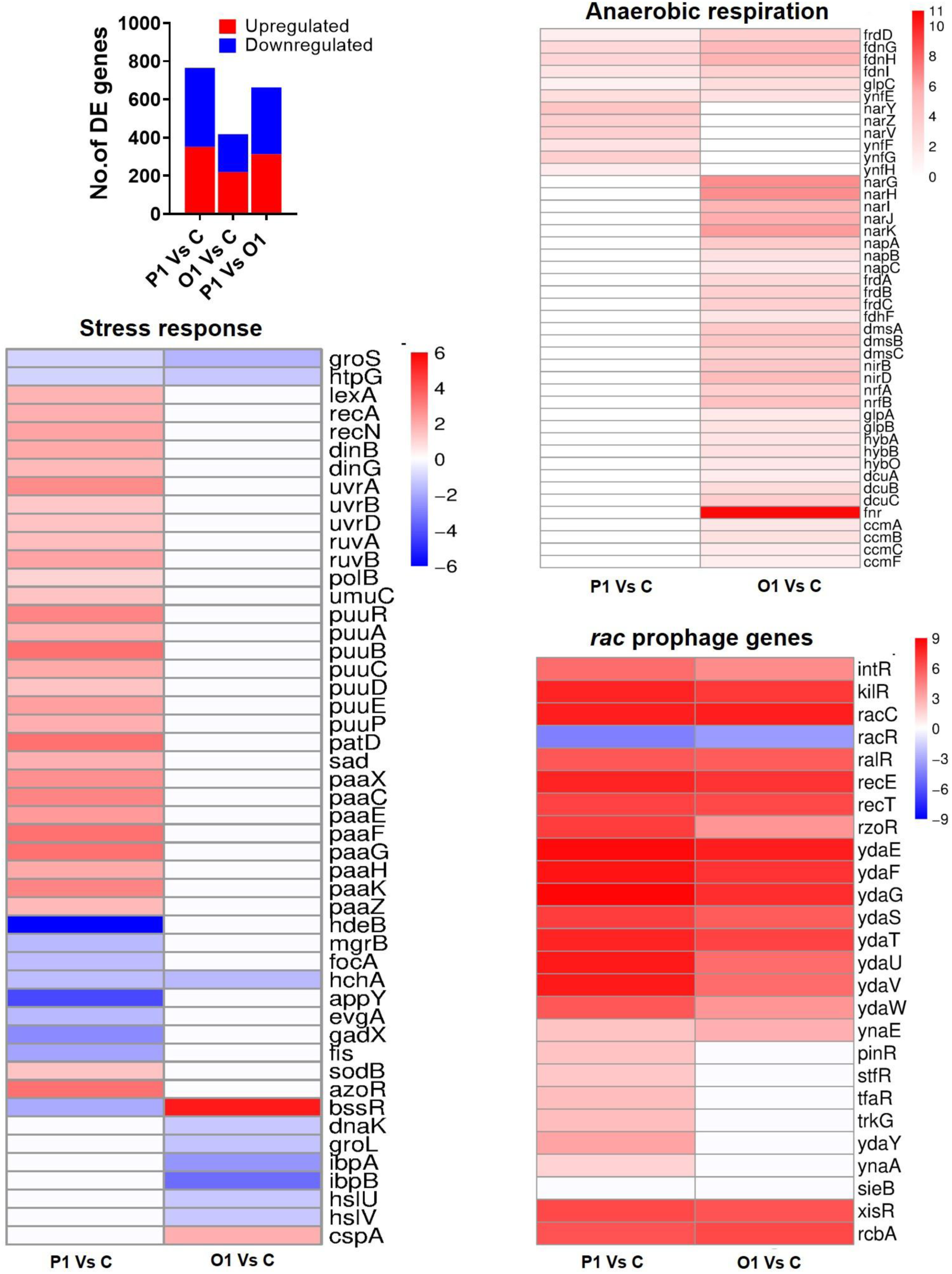
Genome-wide analysis of differential gene expression between the wild type control and *racR* silenced population in *E. coli*. Also see Figure S5-S11, Table S5. Upper left panel: Summary of statistically significant differentially expressed genes, with a bar showing the total number of up and down regulated genes for the tested pairs of strains, (FC ≥ 2, P*adj* < 0.05). Upper right and lower panels: Heatmaps displaying log2 fold transcriptional changes (P1 versus control and O1 versus control) for genes grouped by functional classes: stress response (global stress response and SOS regulon, including DNA-metabolism and repair genes; putrescine and phenylacetate catabolic pathways), anaerobic respiration (core anaerobic respiration, organic-acid and electron transport) and *rac* prophage encoded genes. Colours denote expression changes (red, upregulated; blue, downregulated; white, no significant change; adjusted p ≥ 0.05; non-significant comparisons were set to 0 for visualization). The rows correspond to individual genes. The x-axis represents the log2 FC distribution of genes in each comparison group. The gene order is identical between the P1 and O1 columns to facilitate direct pairwise comparison within each module (no clustering is applied). Exact log2 fold changes and gene identifiers are listed in Table S5: A, D, and E.

Specifically, under stronger silencing, the transcriptional program shifted to an acute stress response. The LexA/SOS regulon was significantly enriched, with induction of genes involved in DNA repair, recombination, and translesion synthesis, including *recA*, *lexA*, *uvrA/B/D*, *recN*, *ruvA/B*, *dinB*, *dinG*, and *umuC*. In parallel, the PspF/phage-shock regulon was strongly enriched, indicating disturbance of inner membrane integrity and proton-motive-force homeostasis (Figure 6: Lower Left panel). Under moderate *racR* silencing, upregulated genes were dominated by regulators of anaerobic and alternative respiration, including NarL, NarP, FNR, DcuR, ModE, RstA, and NikR, together with FlhDC. Consistent with this pattern, genes encoding nitrate and nitrite reductases (*narGHI*, *napAB*, and *nrfA*), fumarate reductase (f*rdABCD*), and nickel uptake (*nikABCDE*) were induced (Figure 6: Upper Right panel). Because the cultures were grown aerobically, this response is most consistent with compensation for impaired energy homeostasis rather than oxygen limitation. Under moderate *racR* silencing, glutamate- and arginine-dependent acid-resistance systems were repressed. These systems are regulated by the Gad network of transcription factors (GadE, GadX, and GadW) and include the structural genes *gadA, gadB, gadE, gadW, gadX*, and the periplasmic chaperones *hdeA, hdeB, and hdeD*. These changes are consistent with withdrawal of energetically costly processes and acid-resistance functions under membrane stress and reduced energy availability. The RacR regulon was among the most strongly enriched sets of genes upregulated in both silencing conditions. The most significant and unique set of regulators directly linked to the physiological effects of RacR depletion is shown in Figure 7. The simultaneous enrichment of the McbR, BluR (YcgE), and H-NS regulons suggests transcriptional reprogramming that integrates biofilm-associated physiology, stress adaptation, and H-NS-mediated regulation of horizontally acquired genomic regions in response to *racR* silencing (47,48). A shared on-target derepression of the Rac cassette and its excision machinery, whereas the secondary response triggered a transcriptional change in the compensatory respiratory program, stress operons, phage shock, and envelope-associated protective systems. These transcriptional changes are consistent with the membrane stress, ATP depletion, and filamentation observed in the accompanying phenotypic assays, indicating that *racR* silencing imposes progressive inner membrane stress.

**Figure 7.**
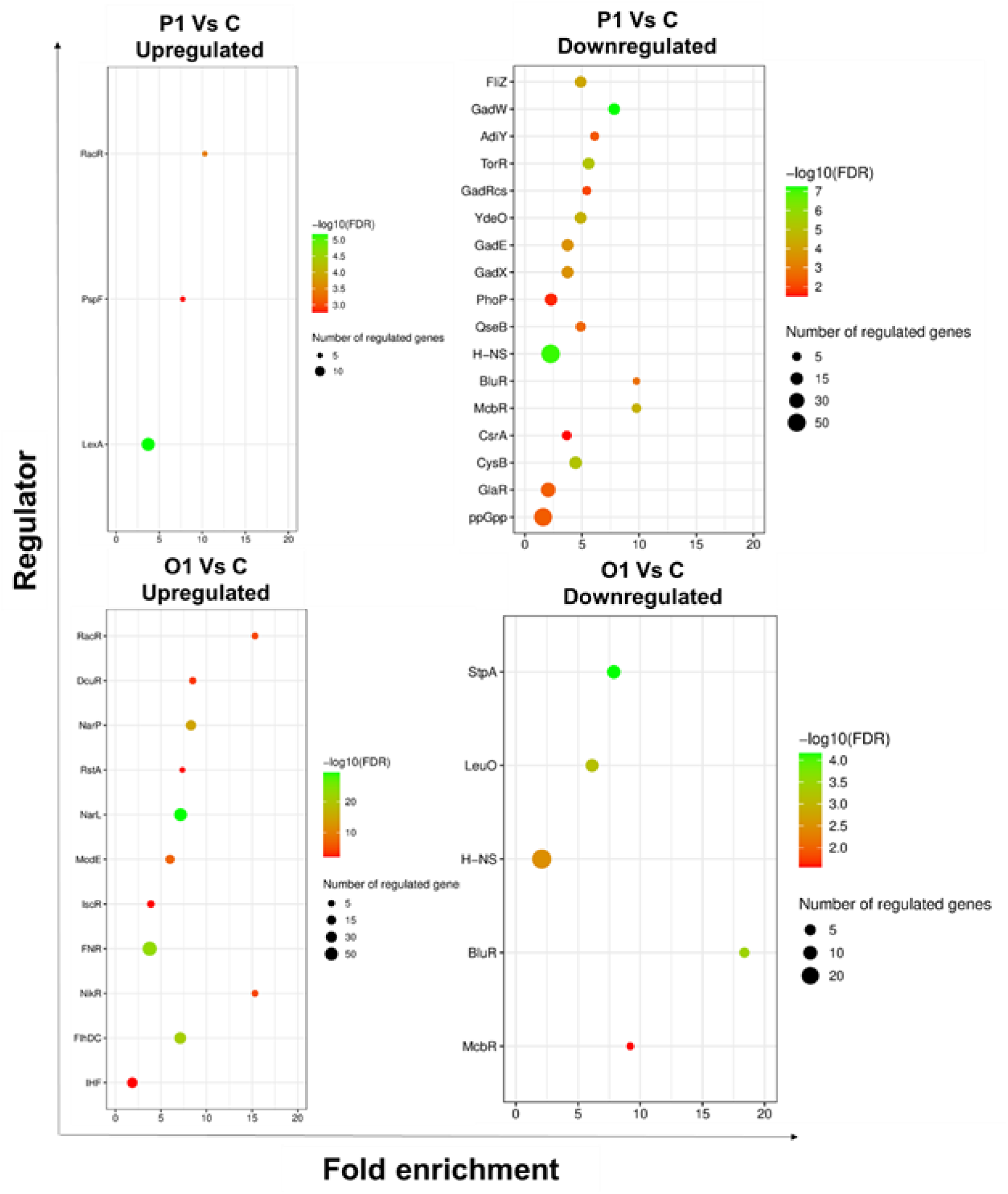
Transcription factor enriched among differentially expressed genes following *racR* silencing. Significantly enriched regulons were identified using a hypergeometric test with Benjamini–Hochberg false discovery rate (FDR) correction (FDR < 0.05 and fold enrichment ≥ 1.5) for both silencing depths (P1 and O1) relative to the control. Each row represents a transcription factor, and the x-axis indicates fold enrichment. The size and colour of the bubble correspond to the number of differentially expressed genes assigned to each regulon and the statistical significance (−log₁₀ FDR), respectively. The RacR regulon served as internal positive control and was significantly enriched under both silencing conditions. Of the 162 significantly enriched regulons identified (listed in Supplementary Data II), 33 regulons with direct relevance to the core physiological and transcriptional responses associated with *racR* silencing are represented.

### 6. Early *rac* prophage excision triggers phenotypic recovery

Previous studies have shown that prolonged exposure to elevated levels of YdaS/T causes Rac excision (16). This was studied either by heterologous expression of C protein or via ectopic expression of YdaT. We asked whether the recovery of stressed cells, as seen in our experimental setup (de-repression of *ydaS/T* induced by CRISPRi knockdown of *racR*), was also due to the loss of Rac from the genome. We performed qPCR on a large number of individual colonies (silenced n= 35; control, n=12) isolated from the induced population at four different time points (3, 5, 8, and 16 h) to detect Rac prophage and study the kinetics of its excision (Figure 8 A).

**Figure 8.**
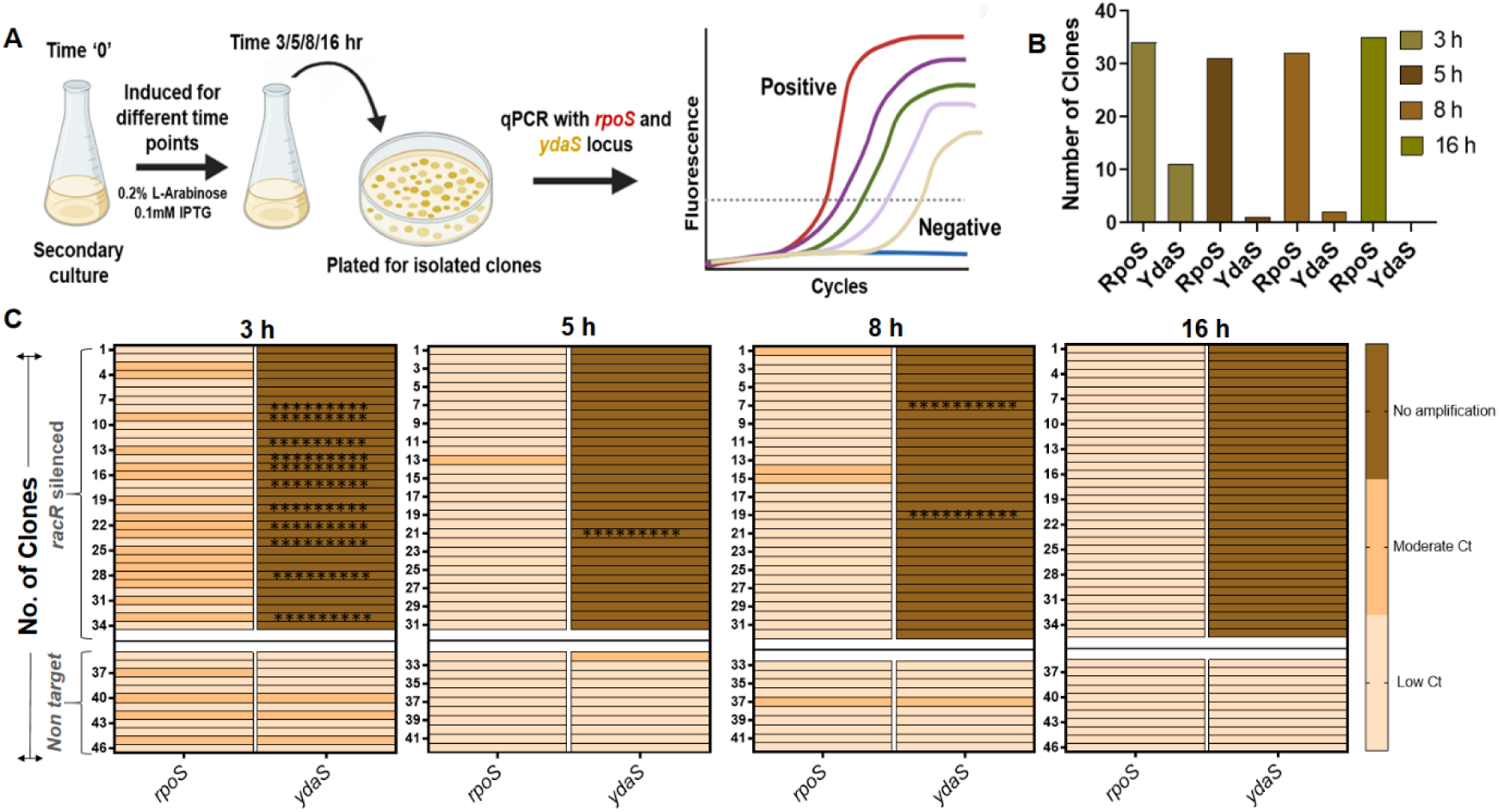
Time dependent loss of specific amplification of *ydaS/T* locus measured by quantitative PCR. (A) Experimental schematic of Rac excision assay. Single colonies were isolated at indicated time points after *racR* silencing, genomic DNA was prepared from individual colonies, and quantitative PCR (qPCR) was performed targeting the *ydaS/T* locus with *rpoS* as an internal reference. (B) Summary of the Rac excision assay showing the total number of clones that yielded positive amplification at different time points. (C) Heat map of Ct values for *ydaS/T* amplification in *racR* silenced (n=35) and non-target control (n=12) clones. Color scale: lower Ct (stronger amplification) = lighter; higher Ct (weaker amplification) = darker. Each row represent an independent colony; column indicates the condition/time point. Asterisks mark colonies from the silenced group that yielded high Ct values but retained a positive melt curve (Tm) consistent with specific amplification. High Ct values without positive Tm were interpreted as non-specific or absent amplification, predominating at 3 h, indicating substantial loss of the ydaS/T template. Complete Ct values, melt-curve calls and amplification metrics are provided in Supplementary data III.

At 3 h post-induction, 11 of the 36 examined clones showed amplification of *ydaS/T* with a positive Tm but very high Ct values. This indicates that only a minority of cells in these colonies carried the Rac prophage at that particular time point. At the same time point, the reference *rpoS* gene was amplified with significantly lower Ct values in all clones, indicating that prophage excision had been initiated. At later time points (5, 8, and 16 h post-induction), only one or two of the total clones yielded detectable Rac-specific amplification, whereas *rpoS* was consistently amplified across all time points with lower Ct values (Figure 8 B & C).

These results reinforce the role of YdaS/T in triggering *rac* excision and its function as a reversible checkpoint that modulates cellular processes, such as replication initiation, cell division timing, and stress metabolism. Our data generated from a larger number of single clones provide much higher resolution and also show that the process occurs at much faster kinetics. qPCR data shows that most of the sampled clones lost RacR within 3 h of induction. The temporal lag observed between DNA loss and morphological recovery reflects the downstream normalisation of the affected cellular processes in the cell cycle. Direct observation of the recovery of viable and morphologically normal cells upon Rac excision provided compelling evidence that the observed phenotype is indeed a reversible, adaptive state that is not associated with permanent cell damage.

### 7. Transient growth arrest with population recovery: Time-lapse microscopy reveals the spectrum of filament fates

To characterize population recovery following the acute toxicity phase of YdaS/T induction, we monitored, cell morphology, membrane potential, intracellular ATP, respiratory activity, and viability at specific intervals within 5-24 h induction period. The cells that continued to grow in the presence of the inducer beyond the 5-hour point exhibited a progressive phenotypic reversal characterized by a reduction in the number of filamentous cells, while normal-morphology rods became increasingly dominant, DiOC₂(3) MFI shifted from the hyperpolarized state to the baseline, and intracellular ATP levels showed partial recovery. By 24 h post-induction, CFU had increased relative to controls confirming that phenotypic reversal was accompanied by a resumption of growth (Figure 9). Recovery can proceed through two entirely different modes which operate exclusively or might co-exist in the population experiencing toxicity. One possibility is autonomous reversal of individual filamentous cells, that is, the same filament that underwent YdaS/T-mediated growth arrest resumes cell division and gives rise to morphologically normal cells. The second possibility is outgrowth competition, in which the filamentous cells are in a terminal or non-recovering state and the reversal is mediated by apparently normal cells, which somehow evaded substantial YdaS/T-driven arrest from the outset. The second possibility also inherently assumes that the filaments eventually undergo lysis. To resolve this, we followed the fate of individual filamentous cells directly using time-lapse microscopy, and investigated whether the filaments were dividing and returning to normal cell state or whether recovery was driven by a subpopulation of cells that remained largely unaffected and proliferated to dominate the population over time.

**Figure 9.**
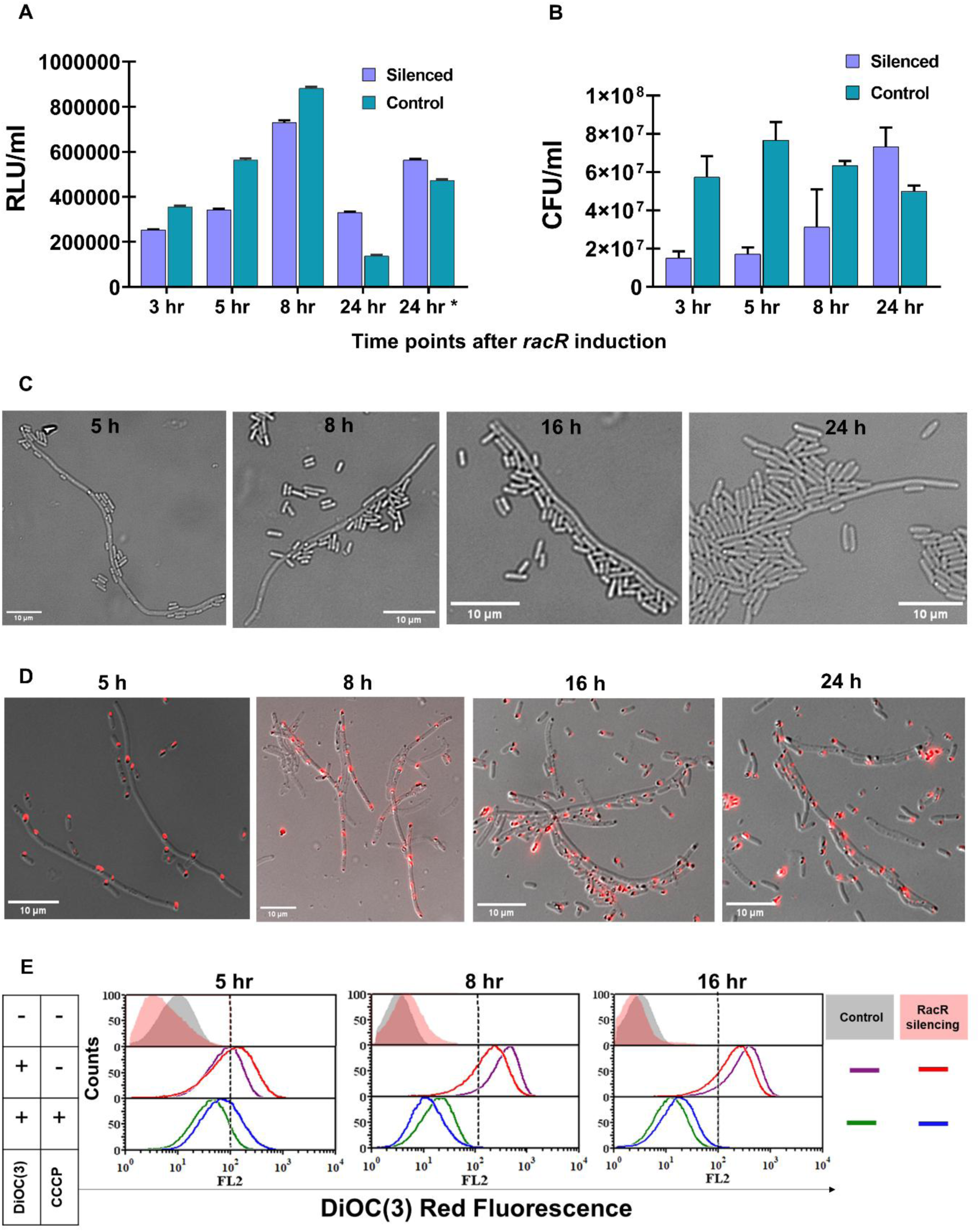
Physiological adaptation following *racR* silencing is accompanied by the recovery of cellular energetics, viability, morphology, and membrane potential. (A) Intracellular ATP levels were measured using the BacTiter-Glo luminescence assay and reported as luminescence (RLU/ml) for both control and silenced cultures over a 24-h period. Initially, ATP levels were notably lower in the silenced cells than in the control cells during the early phase of *racR* silencing, but they gradually returned to normal by 24 h. Transferring 24 h cultures into fresh growth medium restored ATP levels, indicating that the population included metabolically active cells and that ATP deficiency was reversible. The data represent the mean ± SEM of five independent experiments, each performed in triplicate. Statistical analysis was performed using two-way ANOVA (p <0.001). (B) Colony-forming units (CFU) were tracked throughout the time course in *racR* silenced populations. The control group exhibited a decline in CFU at 24 h due to nutrient depletion and entry into the stationary phase, rather than a loss of viability specific to silencing. Data points are presented as mean ± SEM from five independent biological replicates per group. (C) Representative differential interference contrast (DIC) micrographs revealed a mixed population dominated by filaments at early time points, whereas later time points showed a predominance of morphologically normal cells. (D) Representative fluorescence micrographs of CTC (5-cyano-2, 3-ditolyl tetrazolium chloride) staining indicate intracellular formazan deposits in a subset of filamentous cells throughout the time course. (E) DiOC₂(3) staining was analyzed using flow cytometry. Representative histograms demonstrated an increase in red fluorescence in the silenced cells compared to the controls at 5 h, suggesting inner membrane hyperpolarization. This change was negated by the protonophore CCCP, which disrupts membrane potential regardless of silencing status. At later time points, polarization decreased and returned to baseline, relative to the control. The histograms of the control and silenced cells are overlaid across the three experimental conditions, presented alongside the histogram.

Time-lapse microscopy was performed under aerobic conditions. The cells were pre-induced for 2 h, and subjected to time-lapse imaging for 5-6 h. The population displayed a striking range of observations capturing the spectrum of metabolic and physiological heterogeneity (Figure 10). This included relatively shorter filaments that frequently extended into longer filaments (>20 μm), while certain longer filaments underwent elongation and subsequently reverted to rod shapes, with a limited number of instances culminating in lysis during imaging (Movies V1, V2 and V3). There were few instances in which filaments reverted to normal rod-like morphology by synchronous multi-site division events. Additionally, few cells divided by asymmetric division, where new rods were pinched off from one end of the filament (Movies V4, Movie V5 and Supplementary Movie 1). Importantly, the filaments that reverted to rod shapes subsequently divided again, confirming the viability of the newly formed cells (Movie V6). A small subset of filaments exhibited a lemon-shaped central region, commonly referred to as the belly morphology, and maintained this shape throughout the observation period (Movie V7). This indicates localised membrane stress arising either due to inhibited septal PG synthesis or stalled septal constriction (49). A small fraction of filaments spontaneously became translucent during live observation, consistent with cell death without lysis (Movie V8). Conversely, many of the initially longer filaments (∼20 μm) remained growth-arrested throughout imaging, consistent with a persistent, non-recovering state rather than recovery or rapid lysis (Movie V9). Based on these observations, the fate of the filaments could be categorized into three classifications: (1) ‘viable’, if they underwent division at least once; (2) ‘lysed during imaging’, denoting filaments that did not divide and underwent lysis during the image acquisition period; and (3) ‘dead’, which neither divided nor exhibited any increase in length throughout the imaging period or appeared highly translucent.

**Figure 10.**
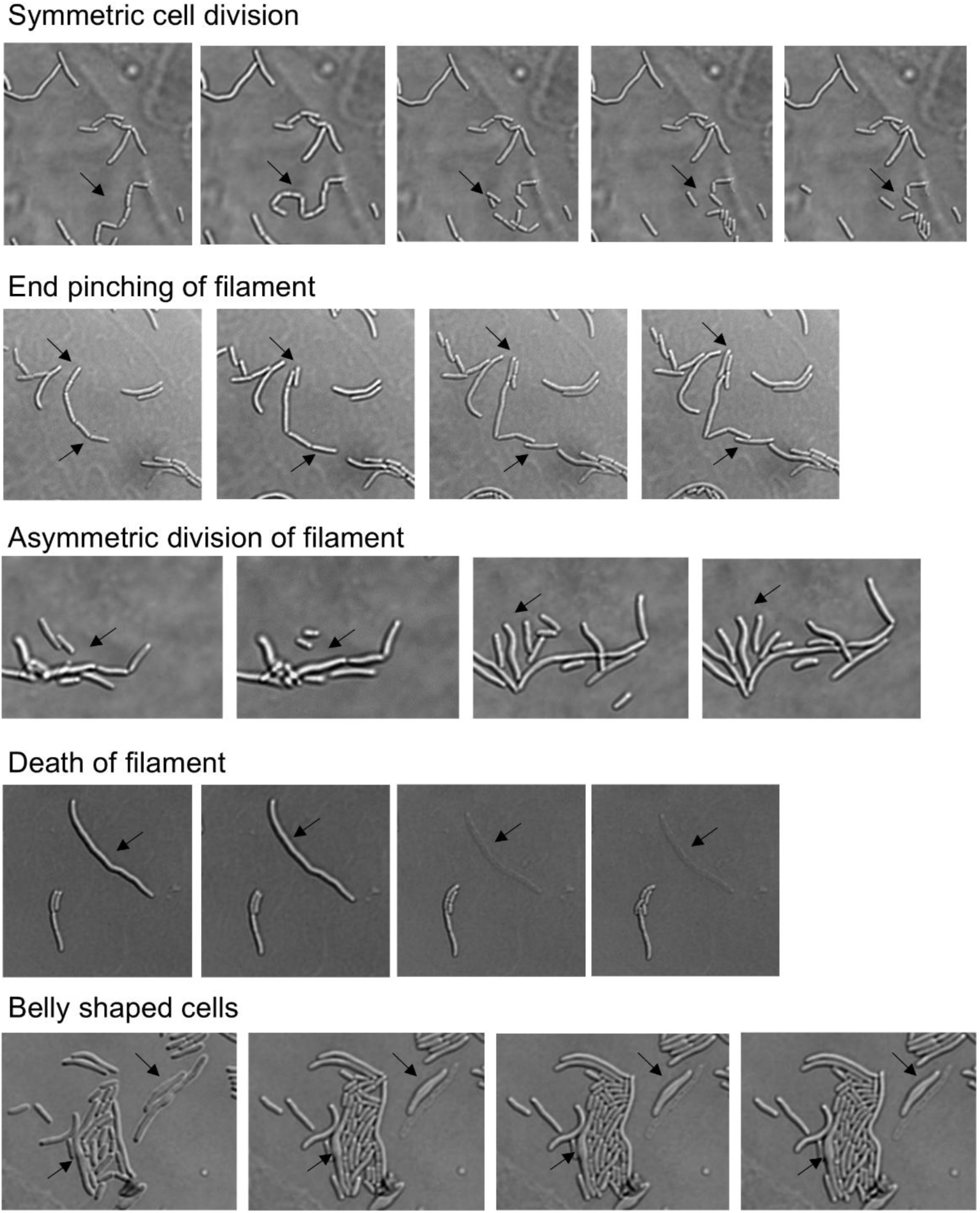
Time-lapse imaging of the filamentous population revealed multiple phenotypic transitions over time. Also see Movie 1-9. Representative montages (shown) were acquired at 60× magnification (scale bar 5 µm) in each panel. The first frame corresponded to t = 0; the same field of view was continuously tracked throughout the observation period. Intermediate frames that exhibited no detectable morphological change are omitted for clarity, with representative observations shown. Sequential panels illustrate discrete events, including (but not limited to) symmetric filament division, asymmetric filament division, cell death, formation of belly shaped cells and end pinching of filaments. Movie 1: Growth of filaments Movie 2: Symmetric division of filament Movie 3: Lysis of filament Movie 4: Asymmetric division of filament Movie 5: Terminal constriction and division Movie 6: Filament-derived cells resumed normal division Movie 7: Belly shaped cells Movie 8: Death of filaments Movie 9: Growth arrested filaments

The transient filamentation may reflects peptidoglycan remodelling, slowed DNA replication, an altered transcriptional program, or a combination of these, which are reset when YdaS/T activity declines after the excision of the prophage. The cells likely engage a controlled metabolic state that temporarily compromises growth while enabling survival during environmental stress, similar to the bet-hedging strategy used by many other bacteria (50).

## Discussion

The Rac prophage has long been recognized as a cryptic element of *E. coli* K-12, losing many structural genes required for productive infection but maintaining a functional but tightly controlled *racR–ydaS–ydaT* regulatory module. Recent literature has drawn an analogy between this module and the well-studied CI-CII-Cro immunity region of lambda, alluding to the lambdoid architecture of Rac where RacR (CI analog) and YdaT (CII analog) cooperation, maintains the prophage in silent mode (51), while RacR depletion and consequent activation of YdaS (Cro analog), causing Rac excision (15). Despite these similarities, there are many differences between these regulatory elements, and their activation has distinct fates for individual cells and the entire population. Rac excision, which does not end in cell lysis, has slower kinetics than λ phage excision. Depletion of RacR, the master regulator of the *racR-ydaST* module, has severe consequences for host cells, including failure of cell division, gross morphological changes, and cell death. The toxic effects are mainly mediated through YdaT, the expression of which is tightly controlled by RacR (12, 17). Interestingly, at the population level, cells eventually recover and resume normal growth. The orchestrated sequence of events following RacR dysregulation, such as the coordinated elevation of YdaT-YdaS levels, YdaT-mediated activation of the Rsc regulon, impaired cell division, and the eventual loss of RacR from host cells, is relatively well documented (16,17). However, a mechanistic understanding of why and how cells cope with lethal stress and the survival strategies employed before lethal stress is relieved by losing Rac is poorly understood. In this study, we aimed to investigate the physiology of cells recovering from the lethal stress imposed by RacR depletion and connect the fates of individual cells to observable phenotypes at the population level.

The most direct physiological signature of reduced RacR expression was selective perturbation of the inner membrane. RacR-silenced cells exhibited inner membrane hyperpolarization, increased EtBr and PI entry, and reduced intracellular ATP with extracellular ATP accumulation, indicating a permeable inner membrane. Moreover, we showed that the filaments were metabolically active and that red fluorescent crystals of reduced CTC were heterogeneously distributed and did not change with time, reinforcing the maintenance of a heterogeneous metabolic state (Figure 9). Simultaneously, the outer membrane barrier remained substantially intact, as evidenced by the lack of growth rescue upon Mg²⁺ supplementation, vancomycin insensitivity, and no significant change in the amount of LPS (Figure 4). In gram-negative bacteria the outer membrane primarily protects against extracellular stress and serves as a selective permeability barrier, whereas the inner membrane is the principal site of cellular metabolism, cell division, solute transport, and proton motive force maintenance. Consequently, selective perturbation of the inner membrane can profoundly impair bacterial physiology, even in the absence of overt disruption of the outer membrane (52, 53). Our results suggest that YdaS/T-mediated toxicity is confined to the inner membrane and is distinct from classical membrane effectors, such as colicins, which affect both membranes. The best characterized inner membrane toxins of *E. coli*, SOS-induced TisB and stringent-response induced HokB, depolarize the membrane through ion-conducting pores (42, 54). In contrast, the hyperpolarization observed in our system suggests restricted proton re-entry and the accumulation of more negative ions in the inner cell environment, excluding pore-mediated collapse.

Such changes in membrane potential are critical because potential is an active requirement for recruiting and anchoring different trans membrane proteins and cell division factors. The amphipathic helix membrane anchors of the division and cell shape proteins, such as MinD, FtsA, and MreB, depend on an intact ΔΨ for correct localization, whereas dissipation or distortion of the potential delocalizes them, blocking septal constriction while leaving the FtsZ ring in place (38,55). Similarly, we observed the recruitment of complete FtsZ rings in a fraction of filaments that failed to constrict and could not complete the downstream steps of cell division. An excessive or dysregulated potential therefore acts downstream of Z-ring assembly, at the recruitment and activation of the late divisome, and the coupling of FtsZ treadmilling to septal peptidoglycan synthesis (56, 57). The multiple, variably spaced rings we observed are the expected readout of a cell that has repeatedly initiated division-site selection while the membrane-potential-dependent constriction step remains blocked. The coexistence of filaments with and without detectable Z-rings suggests marked physiological heterogeneity within the induced cell populations. One possibility is that the concurrent upregulation of the e14 prophage-encoded division inhibitor *ymfM* (sfiC) (36), together with YdaS/T-mediated perturbation of the membrane, prevents the assembly or reassembly of the Z-ring in a subset of filaments, thereby generating filaments without Z-rings. Alternatively, these filaments may represent a terminally compromised subpopulation that has lost the capacity to resume the cell cycle and is ultimately destined for death and lysis. Conversely, filaments retaining Z-rings remain competent to resume cytokinesis upon cellular homeostasis, consistent with the observed reversal of filamentation and bacterial population recovery. Division arrest in our experimental setup is thus the convergence of a perturbed inner membrane affecting late cell division steps and ring assembly by alternative routes, distinguishing the response from other prophage inhibitors such as Rac’s own KilR (58) or Qin prophage DicB (59), which directly targets cell division. However, hyperpolarized, division-arrested cells do not always represent a pathological state. *E. coli* hyperpolarizes as it enters the stationary phase, a poised, low-turnover condition in which growth and division are transiently suspended rather than a state of terminal damage. The arrest imposed by *ydaS/ydaT* derepression mimics this temporary growth arrest, and the cells continue to respire (heterogeneous respiratory activity), retain an intact outer membrane, and halt division without immediately lysing. Similar transient growth arrest has previously been recognized as an adaptive strategy that enhances bacterial survival under stress conditions (60).

Transcriptomics data highlighted the coordinated activation of anaerobic and nitrate-respiratory operons, underscoring the complexity of these inner membrane-mediated physiological effects. The coordinated induction of anaerobic and nitrate-respiratory operons together with the *nikABCDE* nickel-import operon that supplies [NiFe]-hydrogenases represents the respiratory chain remodelling expected of a cell defending its proton-motive force. In addition, alternative terminal reductases are recruited as the balance between proton pumping and re-entry is disturbed (61–63). These transcriptional changes are a compensatory response to emerging bioenergetics loss rather than oxygen limitation under growth conditions. Additionally, the coordinated repression of the Gad acid-resistance regulon supports extensive remodelling of membrane-associated physiology. The glutamate-dependent acid-resistance system consumes cytoplasmic protons during decarboxylation and operates through amino acid antiporters, defending cytoplasmic pH and survival under acid stress, which are functionally coupled to membrane proton homeostasis (64, 65). Cells with compromised inner membrane energetics are unlikely to sustain this PMF-dependent acid resistance system, marking its transcriptional repression as an expected adaptive response to low pH, which is consistent with our physiological assays.

Notably, the transcriptional behaviour of motility genes showed a biphasic response to *racR* silencing. At moderate silencing, the FlhDC regulon and class-2/3 flagellar operons were induced, whereas stronger silencing reversed this pattern and led to the repression of the motility program, marked by the downregulation of FliZ regulon activity (Supplementary Table 5:C and F). This is consistent with an early stress escape response, in which bacteria tend to relocate rather than remodel cellular processes (67, 68). However, as silencing intensifies, YdaT engages the RcsA/RcsB regulon, which in turn represses the FliZ regulon and motility, leading to a stronger stress response. Importantly, these observations reflect a more resolved cellular response to *racR* silencing than what was reported in earlier YdaS/T overexpression studies, which typically resulted in more severe and generalized stress signatures. Our findings complement the YdaT/RcsA regulatory axis characterized previously as YdaT binds directly upstream of *rcsA* and activates its expression, thereby engaging the RcsA/RcsB regulon that governs motility, biofilm formation, colanic acid and capsule synthesis, and cell division (16, 17). Consistent with this model, transcriptomic analysis under our experimental conditions showed the repression of several downstream Rcs-controlled targets, including *csgA*, *fimA,* and flagellar/motility genes, even though *rcsA* and *rcsB* themselves did not meet the differential expression threshold, even under strong silencing. This dissociation is expected as RcsA activity is predominantly controlled post-translation through Lon-mediated proteolysis and H-NS-dependent transcriptional silencing; therefore, a functional increase in RcsA-directed regulation need not be accompanied by a large change in *rcsA* transcript levels (69). However, the RcsA arm accounts for only a part of the toxic phenotype. Gucwa et al. (2024) found that *rcsA* inactivation reduced, but did not abolish, YdaT lethality, explicitly leaving the RcsA-independent component of toxicity unresolved. Therefore, we speculate that membrane bioenergetics perturbation is a strong candidate for the RcsA-independent arm of toxicity, with the relative contributions of Rcs-dependent envelope remodelling and membrane dysfunction likely dictated by expression level, timing, and population heterogeneity. This hypothesis remains to be tested directly in an *rcsA*-null background. Assessment of membrane potential, ATP levels, and permeability in this context would clarify whether the membrane phenotype functions downstream of, in parallel with, or independently of the Rcs pathway.

Time-lapse microscopy revealed divergent single-cell recovery behaviours, underscoring that arrest is reversible but not uniform across the population. The fate of individual cells includes lysis, persistent arrest, and division of individual filaments into viable rods. Direct observation of filament-to-rod reversal is important because bulk recovery could otherwise be explained by the outgrowth of cells that never experienced strong derepression of *ydaS*/*T*. Single-cell tracking has shown that at least part of the population passes through a reversible arrested state and re-enters the cell cycle, consistent with filament recovery by successive divisions after stress (35, 70). Time-lapse microscopy directly demonstrated the reversal of filamentation through both canonical septation and non-canonical division modes, including polar pinching of the filament. Our observation contradicts the assertion of a previous study which states that recovery is mediated by those cells which are resistant to YdaT toxicity (16). We clearly show that filamented cells can revert, undergo division, and thus contribute to population recovery.

The kinetics of Rac excision reported here diverge from those previously described, which we believe is largely attributable to the differences in the experimental setup. Earlier Rac excision was monitored after adventitious repression of *racR* by a heterologous C protein in strains with toxin genes disrupted by *bla* insertions (*ydaS*::*bla* and *ydaT*::*bla*). These manipulations uncoupled *ydaS* and *ydaT* in terms of their native stoichiometry and contribution to excision, while also disrupting the feedback regulation of the intact operon. Excision was quantified at the population level by the fraction of ampicillin-resistant colonies surviving serial passage or by bulk qPCR of total DNA, reporting the accumulation of Rac-excised cells over generations (16). In contrast, our CRISPRi approach perturbs the *racR-ydaS/T* axis in its native regulatory context, allowing toxic and excision programs to operate concurrently, which is a more physiologically relevant scenario. By monitoring excision in individual lineages rather than population averages, we detected Rac excision in more than 70% of individual clones within 3 h indicating early RacR excision in a subset of cells. These results show that Rac excision occurs on a timescale parallel to the onset of YdaS/T-mediated physiological stress, and cell fate is contingent upon the combined outcome of the level, severity, and timing of the toxic effects, as well as the timing of Rac excision. Based on our findings, we propose that *racR* silencing does not elicit a uniform toxic response but instead generates a spectrum of cellular fates, including intact rods, metabolically active filaments, recovering cells, and cells that ultimately lose viability, reflecting heterogeneous membrane dysfunction and metabolic states of the cells. These observations reveal an unexpected morphological plasticity during the recovery process, which has not been previously described in the context of Rac prophage-mediated filamentation. We propose a model (described in detail in Figure 11 and the associated text) that integrates these observations; dysregulation of RacR couples a reversible, inner-membrane-centred division arrest to excision of the element that encodes it, so that fate is set by the race between toxicity and template loss rather than by a single lethal lesion.

**Figure 11.**
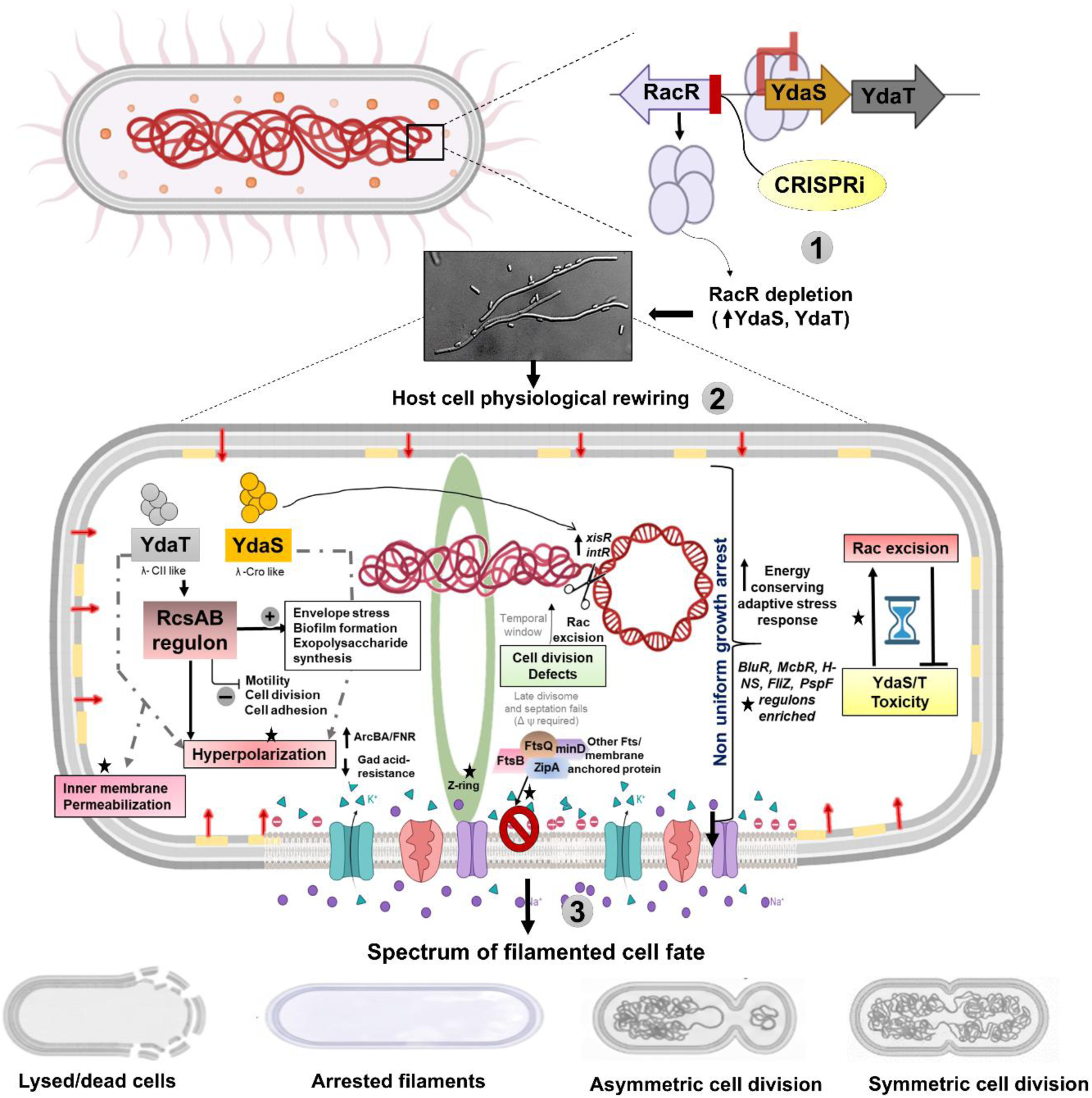
A three-phase model to describing the effects of RacR depletion on cellular physiology and cell fate determination in *E. coli.* (1) In the normal cellular physiological state, RacR maintains lysogeny by repressing the transcription of the prophage-encoded downstream *ydaS/ydaT* locus. CRISPRi-mediated depletion of RacR (λ -CI like) relieves its binding and expresses *ydaS* (λ –Cro like) and *ydaT* (λ -CII like) genes and other genes present on 23kb long Rac prophage. These two soluble effector proteins, YdaS and YdaT induce non-uniform growth arrest and copious filamentation in cells. (2) Activation of the *ydaS/ydaT* toxin locus triggers a cascade of responses centered on inner-membrane energetic defects, including membrane hyperpolarization, ATP depletion, inner membrane permeabilization, and respiratory chain remodelling, leading to transient inhibition of cell division, filamentation, stress-response activation, and Rac prophage excision in a subpopulation of cells. Host responses are mediated by two parallel but interconnected activities: YdaS, which primarily promotes Rac prophage excision, and YdaT, which predominantly drives physiological reprogramming through the RcsA/RcsB envelope stress regulon, with both contributing to energetic and envelope perturbations. Perturbation of membrane homeostasis activates the Psp and ArcBA/FNR systems, inducing an energy-conserving program that suppresses energy-intensive processes, remodels respiratory metabolism, and reinforces envelope stress defenses, reducing proton motive force, stalling Z-ring function, and producing non-uniform, reversible growth arrest; surviving cells eventually restore membrane homeostasis and resume division, and population fate reflects the balance between Rac prophage excision kinetics and the severity of YdaS/T-mediated toxicity. Observations experimentally demonstrated in this study are marked with a star symbol. (3) The spectrum of filamented cell fates following Rac prophage excision and restoration of membrane homeostasis, comprises three outcomes: (i) irreversible lysis and cell death when membrane damage exceeds cellular repair capacity (ii) persistence as dormant filamentous cells when recovery remains incomplete, or (iii) successful recovery into normal, viable rod-shaped cells by symmetric or asymmetric cell division.

In conclusion, we showed that *racR* depletion establishes an integrated, inner membrane centred stress state characterized by selective hyperpolarization, increased inner membrane permeabilization, and an intact outer membrane. We propose that this energetic defect arrests cytokinesis downstream of FtsZ-ring assembly by preventing the recruitment of the late divisome, thereby extending the consequences of RacR loss beyond transcriptional regulation to specific physiological mechanisms. The hyperpolarization-imposed pause holds the filament in a survivable, waiting state until the excision of Rac removes the source of *ydaS/ydaT* expression, permitting the membrane potential to normalize and resume cell division. Cells that undergo extensive membrane permeabilization most likely fail to restore membrane homeostasis and divisome function, ultimately losing viability. Thus, heterogeneous membrane dysfunction and metabolic states appear to determine the balance between recovery and cell death. Transcriptional remodelling corroborates this model, revealing a coordinated energy-conserving response comprising respiratory remodelling, phage shock induction, repression of proton motive force-intensive pathways, and activation of global stress pathways. Collectively, this study provides a mechanistic framework linking prophage activation to bacterial fate. In future, lineage-resolved microscopy combined with *rac* excision reporters should establish the temporal relationship between prophage excision and recovery at the single-cell level, while the genetic separation of YdaS and YdaT contributions will, together, refine the molecular basis of the physiological heterogeneity observed during cryptic prophage activation and, more broadly, how these silent genomic elements shape stress adaptation, persistence, and survival.

## Supplementary Data statement

Supplementary Data are available at NAR Online.

## Resource availability

### Lead contact

Further information and requests for resources and reagents should be directed to and will be fulfilled by the lead contact, Devashish Rath, upon reasonable request.

## Materials availability

No unique reagents generated in this study.

## Data availability

The data supporting the findings of the study are available in this article and its supplementary information files, or from the corresponding author upon request. The raw RNA sequencing data generated in this study are available in the NCBI with BioProject ID PRJNA1503112.

## Acknowledgments

We thank Kavitha Premkumar for assistance with fluorescence-activated cell sorting (FACS) analysis, Celin Acharya and Pallavi S. Chandwadkar for technical support with scanning electron microscopy (SEM), Rahul Checker and Binita for providing access to the fluorescence microscopy, Daniel Daley for generously providing the BS001 strain. The pdCas9 plasmid was a gift from Luciano A. Marraffini and was obtained through Addgene (plasmid #46569).

## Author contributions

Gargi Bindal (Conceptualization, Investigation, Methodology, Formal analysis, Writing – original draft, Writing –review and editing), Devashish Rath (Conceptualization, Supervision, Formal analysis, Project administration, Resources, Writing – original draft, Writing –review and editing)

---

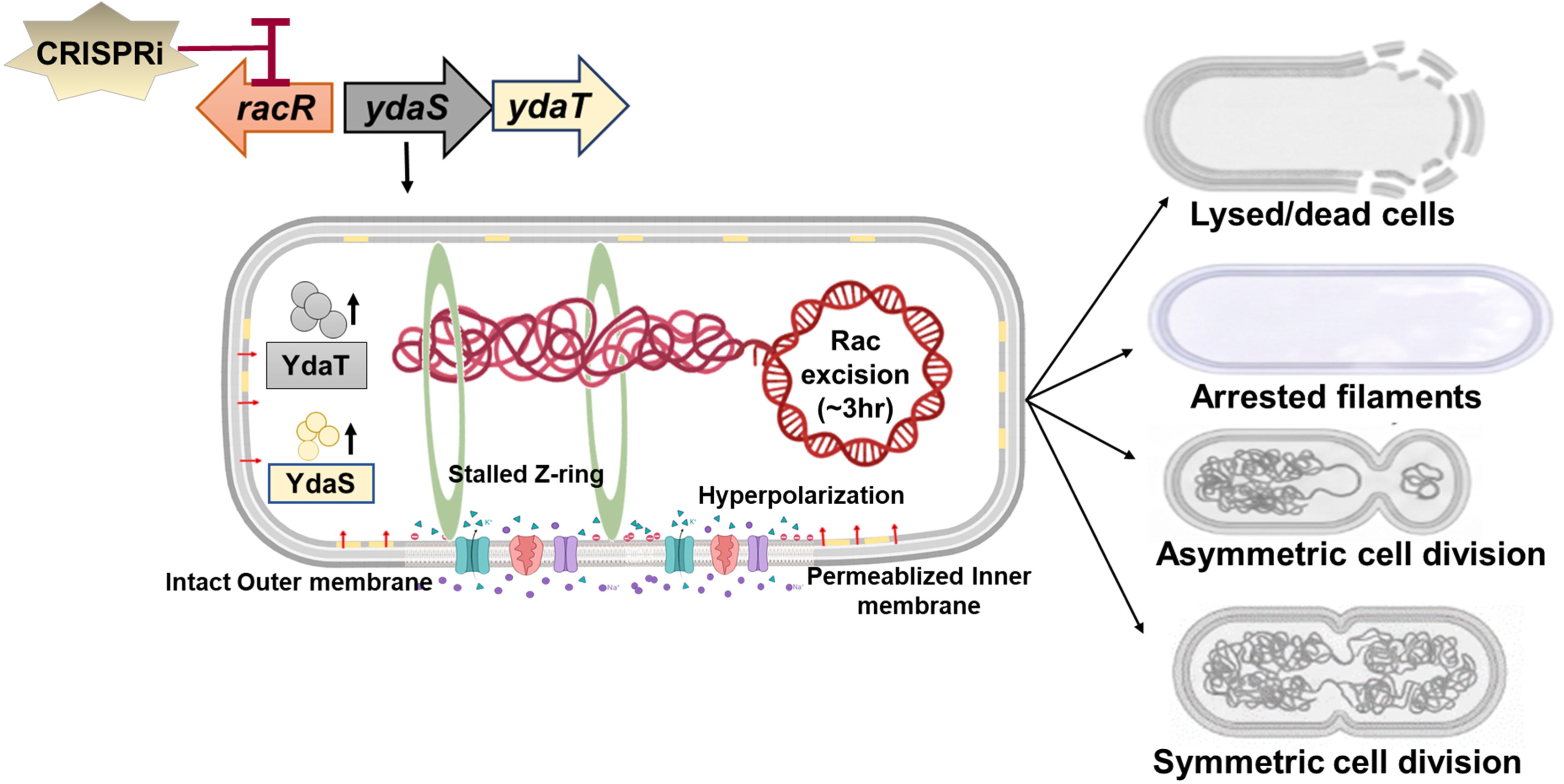

