## Supplementary text file for "CRISPRi-mediated silencing of *racR* reveals molecular strategies for surviving lethal stress coupled to Rac prophage excision"

**Supplementary material**

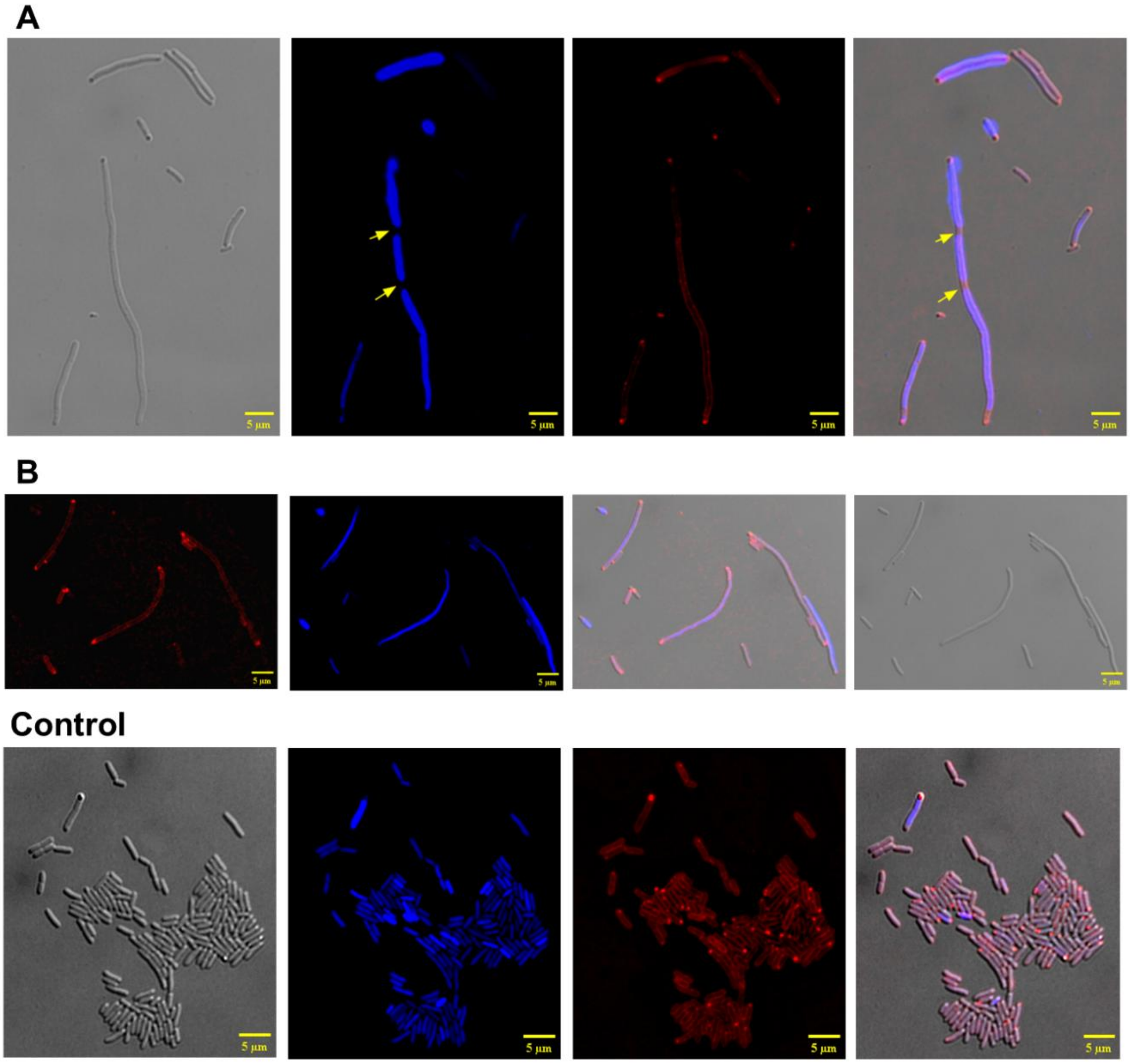


**Figure S1 Fluorescence micrographs of *racR* silenced and control *E. coli* cells stained with DAPI (blue, DNA) and FM4-64 (red, membrane). Related to Figure 1**

(A) Filaments with discrete DNA free gaps between nucleoids (yellow arrowheads), indicating irregular nucleoid segregation or chromosome partitioning defects with intact membrane.

(B) Filaments with continuous nucleoid staining along the cell length with continuous membrane.

(C) Non-targeting control population showing predominantly short rod-shaped cells with regularly spaced nucleoids and uniform membrane staining. Across all panels, FM4-64 staining remains uninterrupted along filaments, demonstrating that elongation occurs without gross membrane structural changes. Images are representative of three independent experiments; for each condition ≥ 300 cells were imaged across biological replicates (Left, Bright field, middle, DAPI and FM4-64; right, merged) (Scale bar 5µm).

.

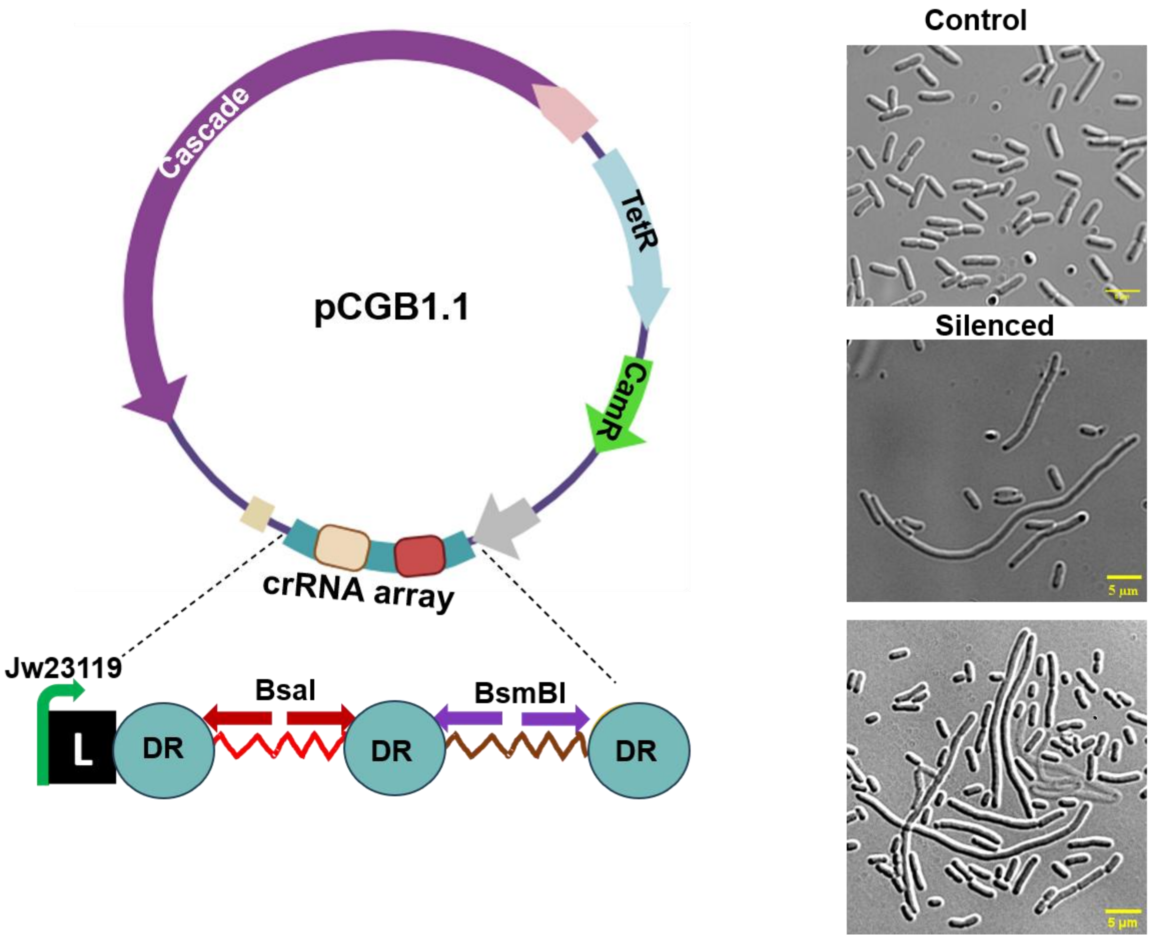


**Figure S2 Construction and validation of an IPTG-independent Type I-E CRISPRi vector for gene silencing in *E. coli*.** Related to Figure 2

(A) Schematic map of all-in-one pCGB1.1 vector. The vector has Cascade under a aTc inducible promoter and a constitutively expressed synthetic crRNA array containing two CRISPR direct repeats flanking the spacer. Target-specific spacers were introduced by a one-step Golden Gate cloning strategy using tandem BsaI and BsmBI Type IIS restriction sites, enabling rapid and directional insertion of annealed oligonucleotides corresponding to new spacer. Representative DIC micrographs showing normal rod-shaped morphology in the non-targeting control and extensive filamentation following silencing of *racR* with pCBG1.1, demonstrating the efficacy of the single-plasmid silencing system. Scale bar, 5 µm.

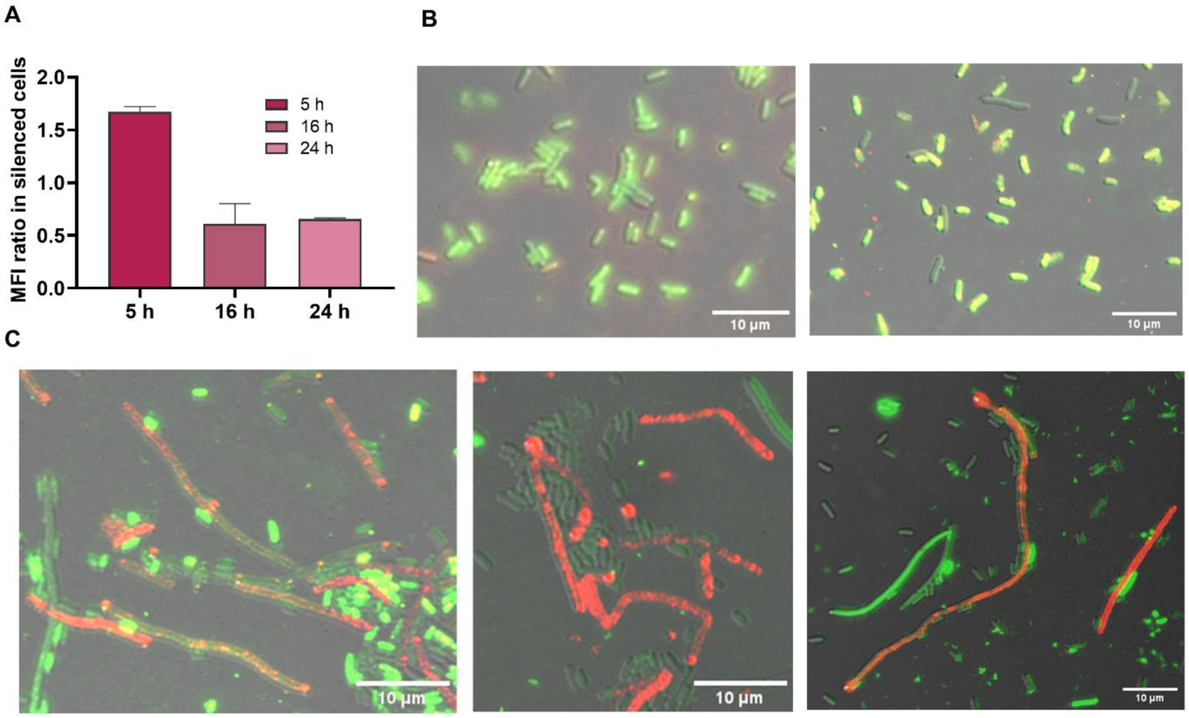


**Figure S3 Time-course and population-level measurement of membrane potential following *racR* silencing**. Related to Figure 3

(A) Time course of membrane hyperpolarization assessed by DiOC2(3) red:green fluorescence ratio in *racR*-silenced and in non-targeting control cells (ratio > 1, hyperpolarized; < 1, depolarized). The MFI ratio is plotted on y axis across indicated time points. *racR* silencing caused increase in the ratio at 5 hour which decreases with time. Mean ± SD, n = 3 biological replicates; two-way ANOVA (p value <0.001)

(B) Population level quantification of membrane potential by multimode fluorescent reader at 5 h post-induction. Mean red:green fluorescence ratio (DiOC2(3)) is shown for *racR*-silenced and control populations (mean ± SEM, n = 10 independent experiments). Statistical significance between groups was assessed by Mann–Whitney U test (p value <0.001).

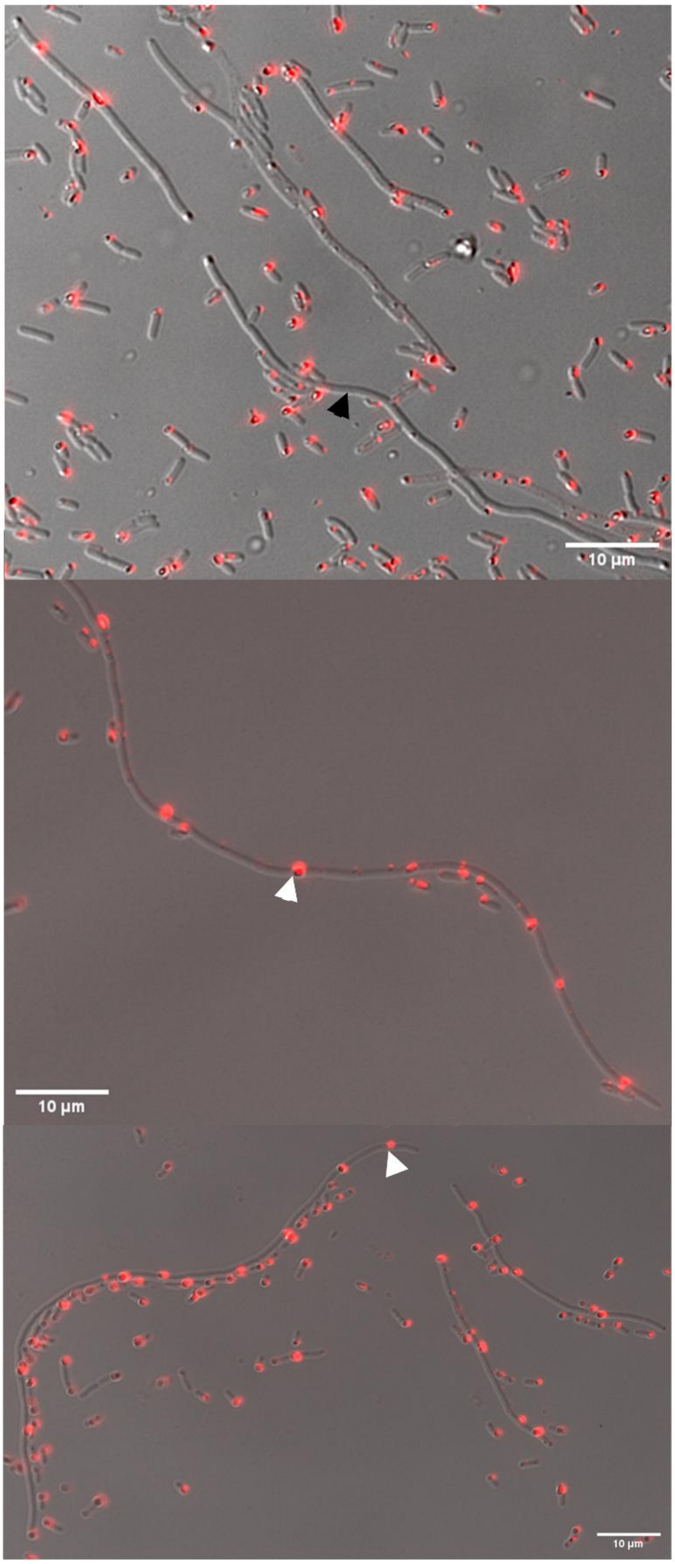


**Figure S4**. **Population level heterogeneity in respiratory activity following *racR* silencing.** Related to Figure 5

Additional representative CTC-stained fields of cells demonstrating cell-to-cell heterogeneity in respiratory activity across the population. White arrows indicate filaments exhibiting CTC formazan deposition, whereas black arrows indicate filaments completely lacking CTC reduction. These additional fields illustrate that respiratory activity is heterogeneous among filamentous cells induced by *racR* silencing. Scale bar, 10 μm.

### Supplementary text section

1. **Global transcriptional changes in presence of *racR* silencing** (related to Results section 5)

To understand the molecular basis of *racR* silencing in host cells, we performed total RNA sequencing for cells carrying *racR* repressing machinery targeted at different positions (P1 and O1) and control cells (C) with non-targeting crRNA. All samples were grown to the exponential phase in minimal medium up to 5 h and total RNA was harvested. The library was prepared using TruSeq Stranded Total RNA kit. The illumina reads produced for the three samples in biological replicates are provided in Table S1. The RNA-seq libraries (P1=3, O1=2, C1=3) generated total of 17.5–26 million raw reads.

**Table S1: Summary of transcriptomics reads and their mapping statistics**.

| Sample_replicate | Reads | Mapped reads | Mapping rate (%) |
|---|---|---|---|
| P1_A | 23022304 | 19976453 | 86.77% |
| P1_B | 25994908 | 23618973 | 90.86% |
| P1_C | 20129374 | 17631319 | 87.59% |
| O1_B | 20787408 | 18099596 | 87.07% |
| O1_C | 17570504 | 15247683 | 86.78% |
| C1_A | 24646722 | 21841925 | 88.62% |
| C1_B | 22659314 | 18555712 | 81.89% |
| C1_C | 17459920 | 13959206 | 79.95% |

Principal component analysis (PCA) was performed using the DESeq2 package (v1.40.2) with biological replicates from the silencing conditions P1 (n = 3), O1 (n = 2), and

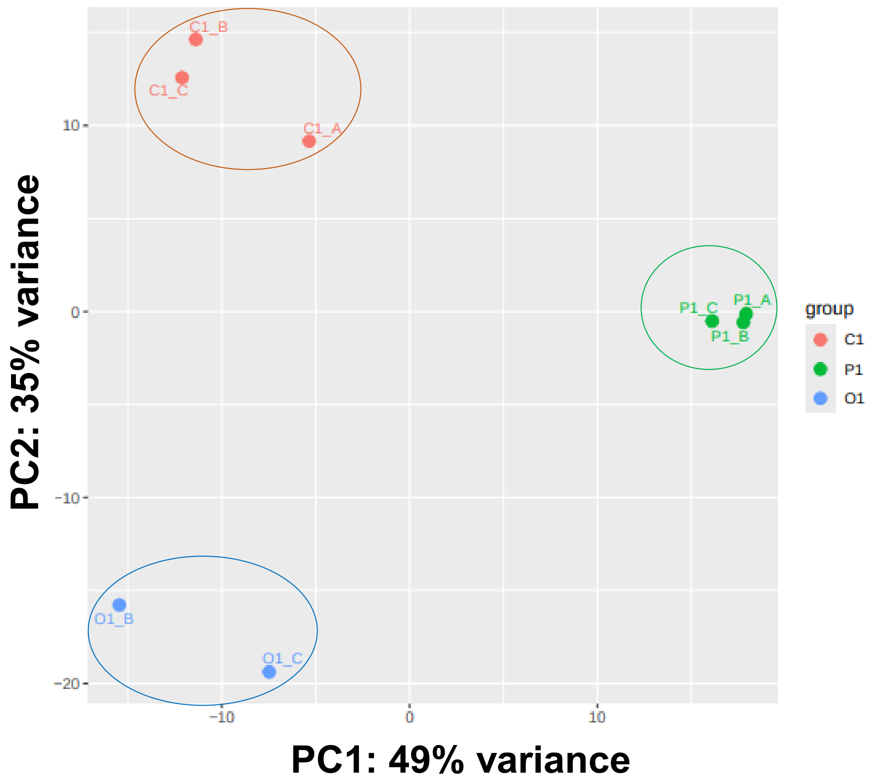


**Figure S5 Principal Component Analysis for the *racR* silenced samples.**

Control C (n = 3) to assess sample clustering prior to differential expression analysis. The analysis showed clear separation between three conditions, while biological replicates were clustered together, which ensures their high concordance per sample (Figure S5). Following quality filtering, 78.03–90.86% of reads mapped to the *E. coli* reference genome, indicating high-quality sequencing data for subsequent transcriptomic analysis. For each pairwise comparison (P1 Vs C, O1 Vs C and P1 Vs O1), differentially expressed genes (DEGs) were defined as those with an adjusted P-value < 0.05 and |log2FC|> 1, corresponding to a minimum two-fold change in gene expression. Among the 4295 quantified genes across all conditions, 765 DEGs in P1 Vs C (350 upregulated and 415 downregulated), 418 DEGs in O1 Vs C (219 upregulated and 199 downregulated), and 662 DEGs in P1 Vs O1 (312 upregulated and 350 downregulated) represented in Figure 6A. Volcano plots were generated using the web tool VolcaNoseR for each pairwise comparison to visualize the overall magnitude of differential gene expression ($log_2$ fold change) (Figure S 6A) (1) . Comparison of the DEGs across the three pairwise comparisons revealed both unique and overlapping transcriptional responses. The shared upregulated genes were predominantly *rac* prophage-associated genes (*intR, rzoR, ydaG, ydaT, ydaU, ydaV, ydaW,* and *ydbJ*), whereas the shared downregulated genes (*cysC, cysH,* and *hdeD*) were associated with sulfur metabolism and acid resistance pathway. The distribution of common and unique DEGs across groups was visualized using Venn diagrams generated with the InteractiVenn web tool (Figure S 6B) (2).

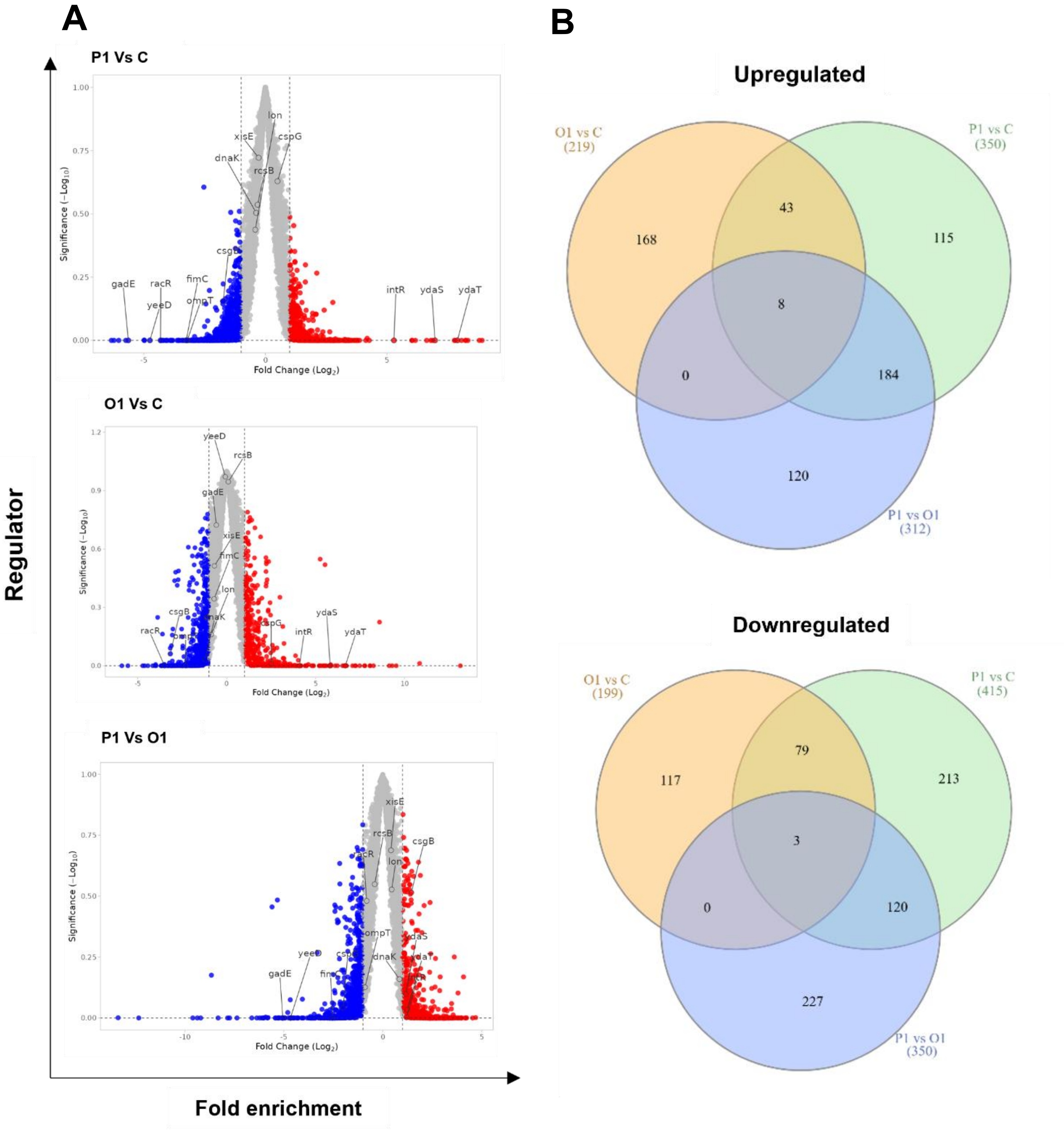


**Figure S6. Differential expression across *racR* silencing conditions.**

(A) Volcano plots represent the overall magnitude of differential gene expression in each pairwise comparison, plotted as log2 fold change versus adjusted significance (Padj $< 0.05$).

(B) Venn diagram showing DEGs unique to or shared among strong (P1), moderate (O1), and control (C) conditions. DEGs were defined as genes with absolute fold change >2 and Padj <

0.05; each sector reports the numbers of up- and down-regulated genes, with total no. of genes in parentheses. See Figure S6 (common genes) and Supplementary data I for full gene lists, fold changes, and statistical values.

Apart from *racR* prophage-associated genes, commonly upregulated genes in P1 and O1 were associated with anaerobic respiration, redox processes, transport, and stress adaptation**.** Heatmap analysis revealed a stronger induction of these metabolic and stress-responsive genes (Figure S7). We also performed qPCR for selected transcripts and the expression trends obtained were consistent with RNA-seq results (Figure S8), confirming the accuracy of transcriptomics profiling.

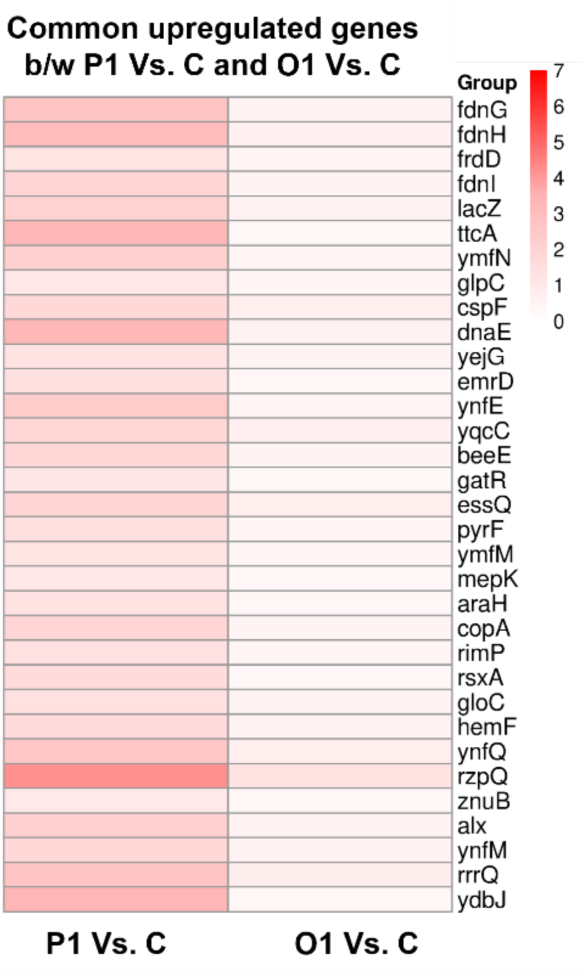


**Figure S7 Heatmap showing the $log_2$ fold changes of common upregulated genes in P1 Vs C and O1 Vs C comparisons (excluding common *racR* genes)**.

Red and blue indicates upregulated and downregulated genes, respectively. Comparisons that were not statistically significant (adjusted $p \geq 0.05$) were assigned a value of 0 for visualization purposes and are shown in white.

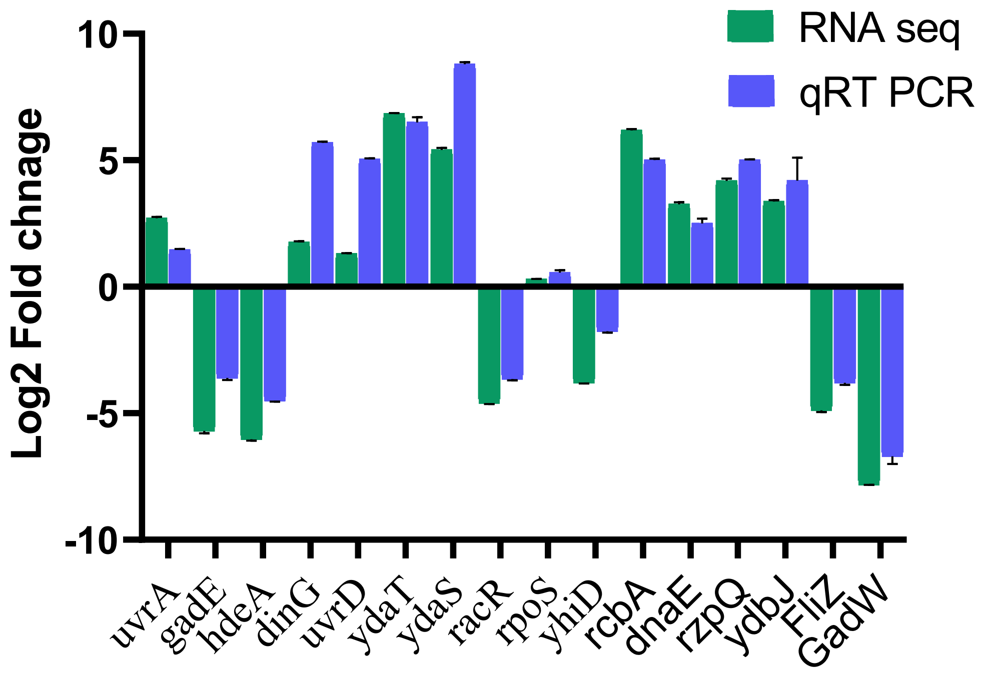


**Figure S8 Validation by qRT-PCR**

Quantitative reverse transcription PCR (qRT-PCR) validation of selected differentially expressed genes identified by RNA-seq under *racR* silencing conditions. Transcript levels were measured in control and *racR*-silenced samples (P1) and normalized to an internal reference gene. The data confirms the direction and magnitude of expression changes observed by RNA-seq, including genes associated with the Rac prophage, stress response, and acid resistance pathways. Relative expression is shown as mean ± SD from three independent biological replicates.

#### 1.1 Functional Enrichment analysis

Functional enrichment analyses were conducted separately for upregulated and downregulated differentially expressed (DE) genes using ShinyGO (v0.85.1), focusing on Gene Ontology categories, Biological Process, Cellular Component, and Molecular Function. An FDR threshold of <0.05 (Benjamini-Hochberg adjustment) was applied to identify significantly enriched pathways, with the entire gene list from each pairwise comparison serving as the background set. These analyses revealed distinct, comparison-specific shifts in cellular functions, highlighting alterations in processes such as respiration, motility, transport, metabolism, DNA repair and recombination, and stress adaptation. For instance, in the P1 versus Control (C) comparison, upregulated genes were predominantly involved in peptide transport, polyamine metabolism, phenylacetate catabolism, DNA repair, and the SOS response, suggesting activation of mechanisms related to cellular maintenance and stress recovery. Conversely, downregulated genes in this comparison were mainly linked to lipopolysaccharide biosynthesis, sulfur metabolism, acid resistance and cell envelope organization, indicating a potential reduction in cell surface structure synthesis and associated metabolic activities (Figure S9). In the O1 versus Control comparison, upregulated genes were enriched in pathways related to anaerobic respiration, electron transport, flagellar motility, chemotaxis, and metal ion metabolism, reflecting enhanced energy production and motility adaptations under altered environmental conditions. Downregulated genes in O1 were associated with stress response and defense mechanisms, implying a possible attenuation of protective pathways (Figure S10). The P1 versus O1 comparison showed upregulation of stress responses, transcriptional regulation, carbohydrate metabolism, peptide transport, DNA repair, and phage defense pathways suggesting a broader activation of cellular maintenance and defense functions. The genes involved in respiratory metabolism, flagellar assembly, transport systems, and lipopolysaccharide biosynthesis are downregulated, highlighted conservative metabolic activity and structural component synthesis (Figure S11). Collectively, these enrichment patterns delineate how gene expression modulations correspond to physiological and adaptive changes across the different experimental conditions.

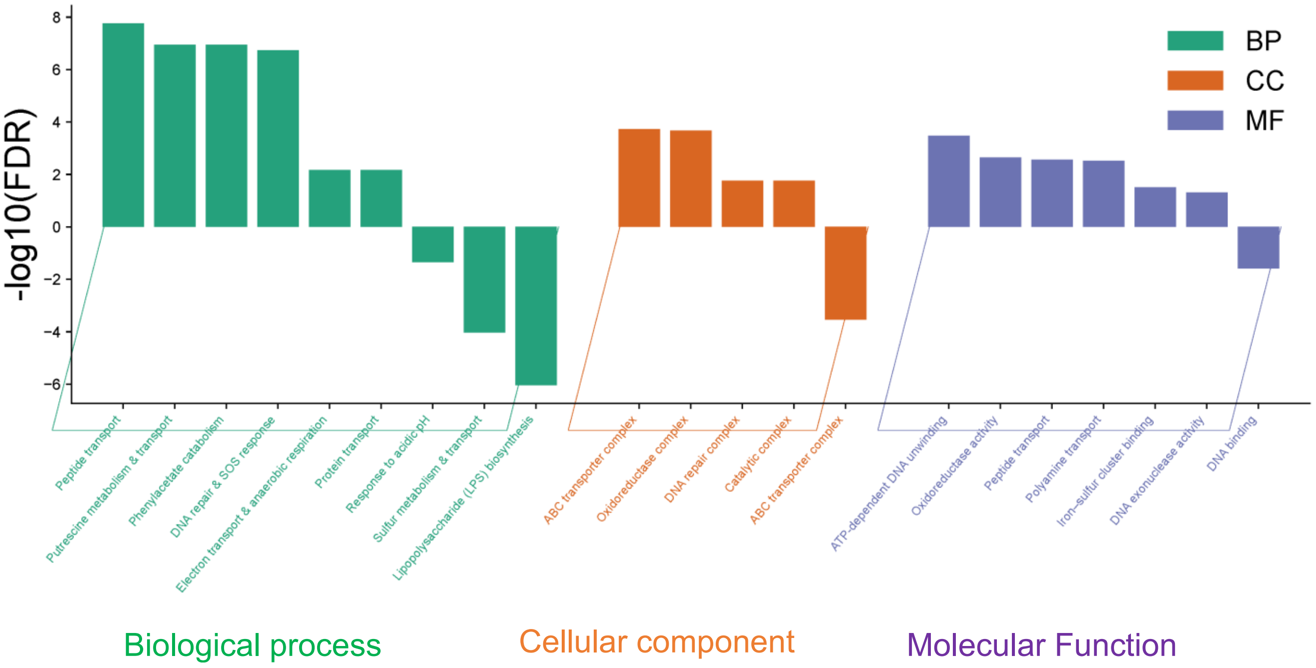

**Figure S9: Gene ontology analysis of the significantly differentially expressed genes for the P1 vs. C comparison.** Enriched GO (BP, CC, and MF) terms are shown as functional clusters. The height of each bar represents the highest $-\log_{10}$(FDR) value among the enriched terms within the corresponding functional cluster. Positive values correspond to enrichment among upregulated genes, whereas negative values correspond to enrichment among downregulated genes. The "biological process" (BP), "cellular component" (CC) and "molecular function" (MF) terms are shown in green, orange, and purple color, respectively

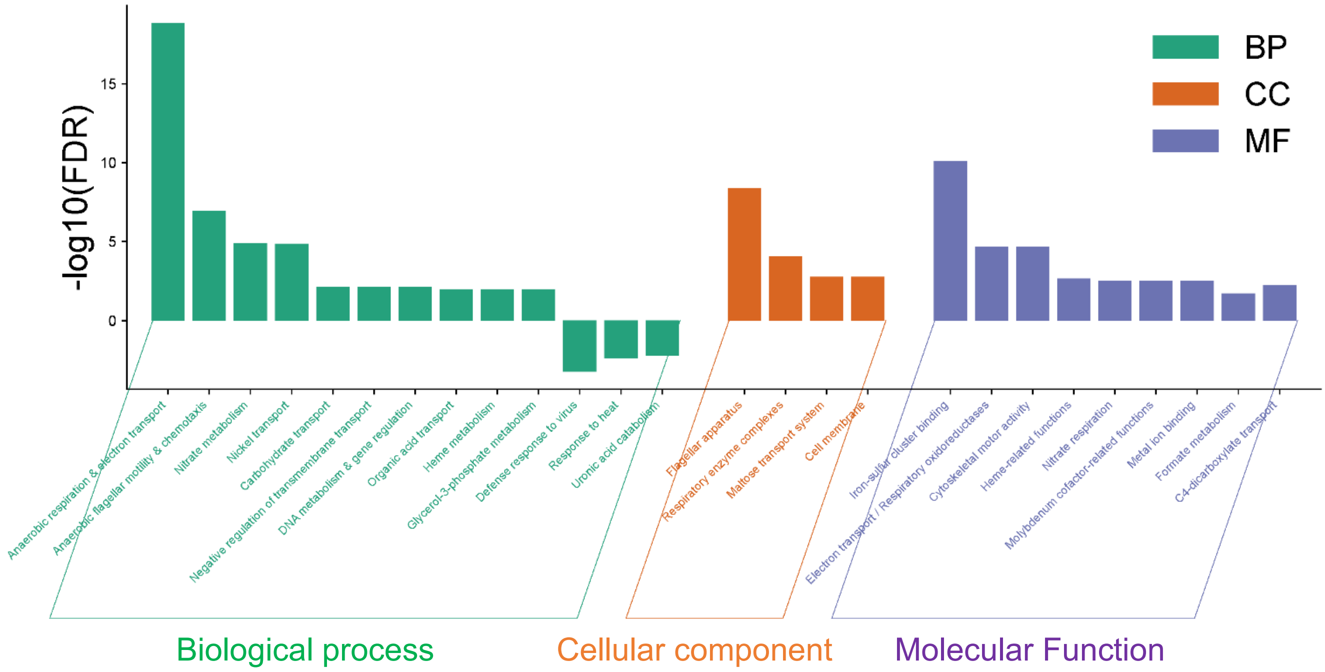


**Figure S10: Gene ontology analysis of the significantly differentially expressed genes for the O1 vs. C comparison.** Enriched GO (BP, CC, and MF) terms are shown as functional clusters. The height of each bar represents the highest $-\log_{10}$(FDR) value among the enriched terms within the corresponding functional cluster. Positive values correspond to enrichment among upregulated genes, whereas negative values correspond to enrichment among downregulated genes. The "biological process" (BP), "cellular component" (CC) and "molecular function" (MF) terms are shown in green, orange, and purple color, respectively.

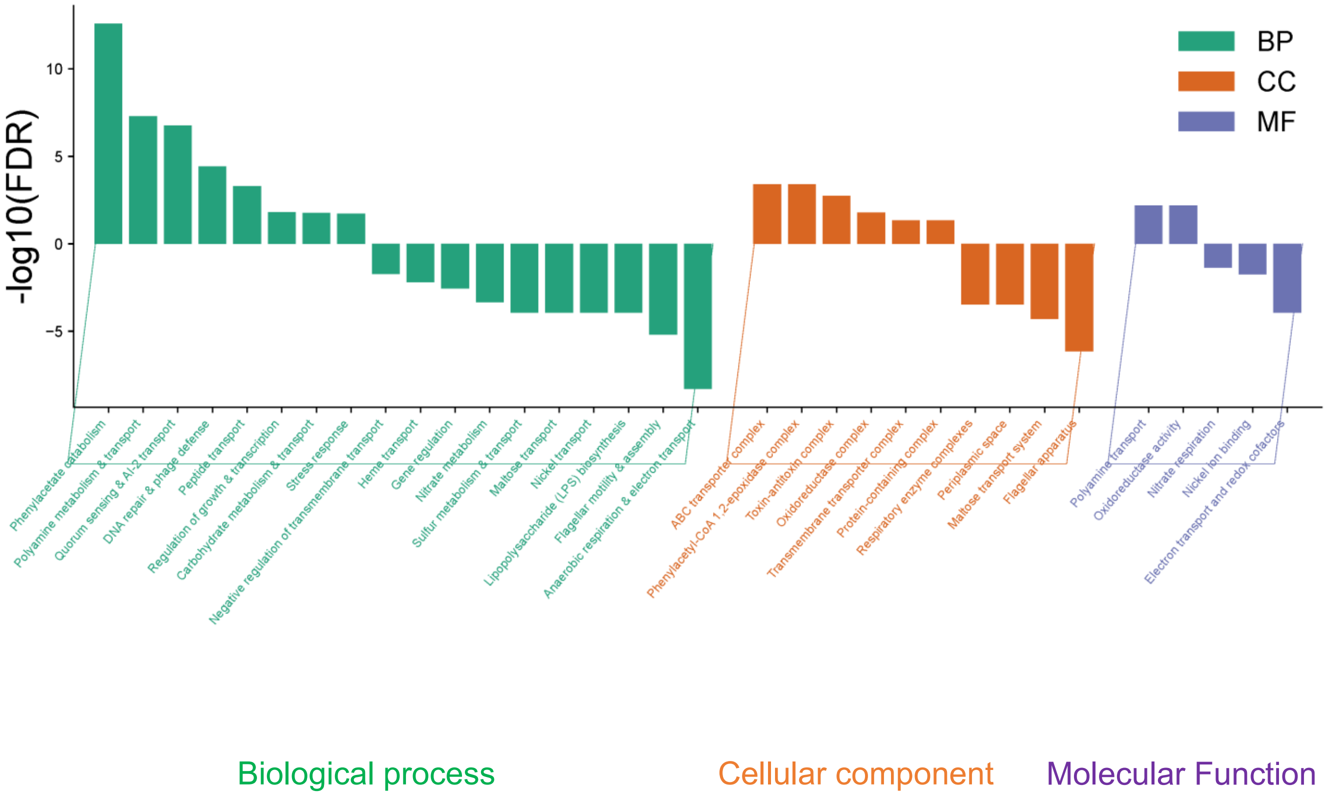


**Figure S11: Gene ontology analysis of the significantly differentially expressed genes for the P1 Vs O1 comparison.** Enriched GO (BP, CC, and MF) terms are shown as functional clusters. The height of each bar represents the highest $-\log_{10}$(FDR) value among the enriched terms within the corresponding functional cluster. Positive values correspond to enrichment among upregulated genes, whereas negative values correspond to enrichment among downregulated genes.The “biological process” (BP), “cellular component” (CC) and “molecular function” (MF) terms are shown in green, orange, and purple color, respectively.

Further, transcriptional regulatory interactions were obtained from the RegulonDB RegulatorGene dataset and analyzed using custom Python script (Figure S12) to identify transcriptional regulators whose experimentally validated target genes overlapped with differentially expressed gene (DEG) sets. For each pairwise comparison and expression direction (upregulated and downregulated), DEG lists were compared against the processed RegulonDB dataset to identify regulators with overlapping DEGs targets.

```
# Convert DEG genes into a searchable set
deg_genes = set(deg["Gene"].astype(str))
# Identify regulators with DEG target overlap
for _, row in regulon.iterrows():
regulator = row["Regulator"]
# Extract experimentally validated target genes
target_genes = set(g.strip() for g in row["Regulated_Genes"].split(","))
# Identify overlapping genes between regulator targets and DEGs
overlap = target_genes.intersection(deg_genes)
# Retain regulators with at least one DEG target
if overlap:
results.append([regulator, overlap])
```

**Figure S12 Schematic workflow illustrating the pipeline used to extract regulator-associated DEGs across *racR* silencing conditions**.

DEGs from each pairwise comparison mapped to RegulonDB-based regulator target sets to identify overlapping and uniquely enriched regulators. The overlap analysis was used to compare condition specific and shared regulatory responses, enabling prioritization of regulators most likely to contribute to the physiological effects of *racR* silencing. Enrichment results were visualized as heat maps (related to Figure 7) and summarized in table to highlight regulators linked to core physiological processes such as envelope stress, respiration, motility, and global stress adaptation affected in *racR* sielncing (complete list of regulators provided in Supplementary Data II).

The identified regulator-DEG overlaps were subsequently used for transcriptional regulator enrichment analysis and visualization. The regulator-DEG overlap analysis identified regulators associated with each differentially expressed gene set. In the P1 vs C comparison, 96 regulators were associated with the upregulated DEGs and 152 with the downregulated DEGs. In the O1 vs C comparison, 85 regulators were associated with the upregulated DEGs and 124 with the downregulated DEGs. For the P1 vs O1 comparison, 109 regulators were associated with the upregulated DEGs and 132 with the downregulated DEGs. These regulator DEG associations formed the input for the subsequent transcriptional regulator enrichment analysis. The enrichment was assessed by the hypergeometric upper-tail test, comparing the overlap between each RegulonDB regulon and the differentially expressed gene set against that expected by chance from a total of 2,793 genes. Fold enrichment was calculated as the observed overlap divided by the expected overlap, and P values were corrected for multiple testing by the Benjamini–Hochberg procedure. Regulons with FDR < 0.05 and fold enrichment

≥ 1.5 were considered significantly enriched and are listed in Supplementary Data II. Pathway enrichment bar plots, heatmaps, and bubble plots were generated using the web-based platform SRplot (3).

**Table S2: Bacterial strains used in this study.**

| Strain name | Genotype | Remarks | Source |
|---|---|---|---|
| GB051 | MG1655, Δ*cas3*, *araB*::T7RNAP-tetA, *racR*:: 3XFLAG | Used to conduct all experiments in this study, unless otherwise indicated | (6) |
| MG1655::FtsZ | MG1655, Δ*cas3*, *lacI*::*ftsZ–gfp* | Expresses FtsZ-GFP under 2.5 µm IPTG | This study |
| BS001 | MG1655 (*lacI*$^{q}$ *P*$_{208}$*- ftsZ –gfp)* | Used as source of FtsZ-GFP | (7) |

**Table S3: Plasmids used in this study.**

| Plasmids | Remarks | Source |
|---|---|---|
| pWUR400 | Kind gift | (4) |
| pdCas9 | Kind gift | (5) |
| pZE12luc-P1 | pZE12luc expressing crRNA with P1 spacer | (6) |
| pZE12luc-O1 | pZE12luc expressing crRNA with O1 spacer | (6) |
| pZE12luc-O2 | pZE12luc expressing crRNA with O2 spacer | (6) |
| pZE12luc-NT | pZE12luc expressing crRNA with NT spacer | (6) |
| pCBG1.1 | aTc-inducible all-in-one Type I-E CRISPRi vector for *racR* knockdown | This study |

**Table S4: Oligonucleotides used in this study.**

| **Oligonucleotides** | **Sequence** | **Remarks** |
|---|---|---|
| *ydaS* Fwd | TGTGCTGTTGTCGGTGGGCA | qRT-PCR |
| *ydaS* Rev | GCTGGACATCTCTCGGCAGGC | qRT-PCR |
| *ydaT* Fwd | CTTCGATACCCTGGAACGCC | qRT-PCR |
| *ydaT* Rev | AGAATCGTCGGAAGAACCGC | qRT-PCR |
| *racR* Fwd | TCAAAGGCGGAGGTCGCACG | qRT-PCR |
| *racR* Rev | GCCCCCAATGCTCTGGACCAA | qRT-PCR |
| MG1655 16S RNA Fwd | AGCCTGATGCAGCCATGCCG | qRT-PCR |
| MG1655 16 S RNA Rev | AGCCGGTGCTTCTTCTGCGG | qRT-PCR |
| *uvrA* Fwd | CAAATCCTCGCTCGCTTTCG | qRT-PCR |
| *uvrA* Rev | GCAGGAGAAAGCCCCTCAAT | qRT-PCR |
| *gadE* Fwd | TGGAGAAATTAGATGCCGAGAG | qRT-PCR |
| *gadE* Rev | TTGATACTTTCTTTGCGGCTAAC | qRT-PCR |
| *hdeA* Fwd | CTGAAGCGCTGAACAACAAAG | qRT-PCR |
| *dinG* Fwd | ACTGATTCGAAGCCACGGTT | qRT-PCR |
| *dinG* Rev | CATCCAGTAGTCGCTTGCCA | qRT-PCR |
| *uvrD* Fwd | CTACGGTAATCCGGTGGAGC | qRT-PCR |
| *uvrD* Rev | GTTGCAGGATATGCGGCTTG | qRT-PCR |
| *yhiD* Fwd | CGGTGCAGGTAACATTCTCG | qRT-PCR |
| *yhiD* Rev | CCGCTGCCAATAACCATACC | qRT-PCR |
| *rcbA* Fwd | GGTGGCACTCCTACTAACTGG | qRT-PCR |
| *rcbA* Rev | TCTGGAAACGCCTGCTTCTT | qRT-PCR |
| *dnaE* Fwd | CGATCAAAGGGGTCGGTGAA | qRT-PCR |
| *dnaE* Rev | CACGCGACGGTTCAACTTTT | qRT-PCR |
| *ydbJ* Fwd | AATTGTGCAATGATCGGCGG | qRT-PCR |
| *ydbJ* Rev | TGACTGTTCGCTACAGCGTT | qRT-PCR |
| *FliZ* Fwd | AAATGGCTGGCAAACGGAAC | qRT-PCR |
| *FliZ* Rev | CACCGTACCTGGGCTCATTT | qRT-PCR |
| *GadW* Fwd | TTTGTTCACCGGATACGCGA | qRT-PCR |
| *GadW* Rev | TTGCGTGGTAGCTGACGAAT | qRT-PCR |
| *ydaS* Fwd | TGTGCTGTTGTCGGTGGGCA | Rac excision assay |
| *ydaT* Rev | AGAATCGTCGGAAGAACCGC | Rac excision assay |
| *rpoS* Fwd | GGTAAAAATTGCCCGCCGTT | Rac excision assay |
| *rpoS* Rev | TGAGAAGCGGAAACCACGTT | Rac excision assay |
| Cascade_Tet_ Fwd | TCAGCTAGGAGGTGACTAACATGGCTAATTTGCTTATTGATAACTG | Amplification of Cascade |
| Cascade_Tet_Rev | TAGATCCTTTCTCCTCTTTATCACAGTGGAGCCAAAGATAG | Amplification of Cascade |

**Table S5 Differential expression of functionally related gene groups following racR silencing.**

| A. Coordinated expression of the Rac prophage genes upon *racR* silencing | | | | |
|---|---|---|---|---|
| Gene | Locus tag | log2(FC) P1 Vs C | log2(FC) O1 Vs C | Function |
| *racR* | b1349 | -4.31314110551252 | -3.49843800030548 | master repressor (**CRISPRi target**) |
| *racC* | b1351 | 8.06774012294212 | 8.06199837975548 | Rac lysogeny/structural |
| *ydaS* | b1341 | 6.97947280878199 | 5.81828654831153 | Putative toxin |
| *ydaT* | b1342 | 7.89991697617435 | 6.66966325179903 | Putative toxin |
| *ydaE* | b1348 | 8.80248084697272 | 8.05768118189718 | Rac prophage protein |
| *ydaF* | b1347 | 8.3314483372861 | 7.22630046820812 | Rac prophage protein |
| *ydaG* | b1346 | 8.91345232012826 | 7.47586210428566 | Rac prophage protein |
| *ydaU* | b1345 | 8.23290198852269 | 5.22728203852638 | Rac prophage protein |
| *ydaV* | b1344 | 8.24631663402199 | 5.21161652680262 | Replication protein |
| *ydaW* | b1343 | 6.03559539886122 | 3.75183783698955 | Rac prophage protein |
| *recE* | b1350 | 7.80297859422681 | 7.24326765955462 | Exonuclease VIII |
| *recT* | b1349 | 6.80865939434287 | 6.51757397861774 | Recombinase |
| *kilR* | b1330 | 7.7779380662043 | 7.07590456948113 | Cell division inhibitor |
| *ralR* | b1352 | 6.02686792462948 | 5.778715553748 | Toxin |
| *xisR* | b1356 | 6.63513977932231 | 6.09812626306829 | Excisionase |
| *intR* | b1357 | 5.28820735779302 | 4.09189982188406 | Integrase |
| *rzoR* | b1358 | 6.87917136565898 | 3.83719681601806 | Lysis protein |
| *rcbA* | b1356 | 6.17088715999537 | 6.60230280779042 | Rac-associated |
| ynaE | b1375 | 2.14535886789916 | 2.82417583198268 | Cell division regulation |
| pinR | b1374 | 2.17087986969464 | NS | DNA inversion |

| | | | | |
|---|---|---|---|---|
| tfaR | b1373 | 2.30476955761622 | NS | Phage transcription activation |
| stfR | b1372 | 1.92233854797893 | NS | Tail fiber protein |
| trkG | b1363 | 2.30352650460795 | NS | Potassium transport |
| ydaY | b1366 | 3.34305601514497 | NS | Rac prophage protein |
| ynaA | b1368 | 1.69383343628407 | NS | Stress response |
| sieB | b1353 | NS | NS | Super infection exclusion |

**B. Common upregulated genes among P1 Vs C and O1 Vs C (excluding *racR* genes)**

| Gene | Locus tag | log2(FC) P1 Vs C | log2(FC) O1 Vs C | Function |
|---|---|---|---|---|
| *xisR* | b1346 | 6.63514 | 0.559077 | Phage excisionase involved in rac prophage DNA recombination |
| *rcbA* | b1347 | 6.170887 | 0.686296 | Rac prophage protein; Involved in DNA double-strand break suppression and genome stability |
| *fdnG* | b1474 | 2.742261 | 0.624731 | Involved in anaerobic respiration |
| *fdnH* | b1475 | 3.057167 | 0.694885 | Involved in anaerobic respiration |
| *frdD* | b4151 | 1.245562 | 0.452755 | Membrane anchor subunit of fumarate reductase; Involved in anaerobic respiration |
| *fdnI* | b1476 | 1.918974 | 0.498437 | Involved in electron transfer |

| | | | | |
|---|---|---|---|---|
| *lacZ* | b0344 | 2.062479 | 0.592334 | β-galactosidase involved in lactose metabolism |
| *ttcA* | b1344 | 3.272731 | 0.357464 | Involved in tRNA modification |
| *ymfN* | b1149 | 2.127799 | 0.483971 | Rac prophage protein; Involved in DNA damage response and genome maintenance |
| *glpC* | b2243 | 1.075167 | 0.442805 | Iron-sulfur electron transfer subunit of anaerobic glycerol-3-phosphate dehydrogenase |
| *cspF* | b1558 | 1.807246 | 0.736139 | Cold shock RNA-binding protein involved in stress adaptation |
| *dnaE* | b0184 | 3.259065 | 0.576935 | Involved in chromosomal DNA replication |
| *yejG* | b2181 | 1.302995 | 0.50646 | Uncharacterized conserved protein of unknown function |
| *emrD* | b3673 | 1.418358 | 0.411152 | Multidrug efflux transporter involved in export of toxic compounds |
| *ynfE* | b1587 | 2.368334 | 0.479863 | Iron-sulfur electron transfer protein involved in anaerobic oxyanion reduction |

| | | | | |
|---|---|---|---|---|
| *yqcC* | b2792 | 1.869224 | 0.660743 | Uncharacterized conserved protein of *E. coli* |
| *beeE* | b1151 | 1.897677 | 0.5159 | Putative bacteriocin-associated protein involved in bacterial defense |
| *gatR* | b4498 | 1.157023 | 0.35174 | Transcriptional regulator controlling galactitol utilization |
| *essQ* | b1556 | 1.926148 | 0.774592 | unknown function |
| *pyrF* | b1281 | 1.415042 | 0.449947 | Involved in pyrimidine biosynthesis |
| *mepK* | b0926 | 1.129013 | 0.370364 | Involved in bacterial cell wall remodeling |
| *araH* | b4460 | 1.27464 | 0.393183 | Membrane component of the L-arabinose ABC transporter involved in arabinose uptake |
| *copA* | b0484 | 1.974493 | 0.530395 | Copper-exporting P-type ATPase involved in heavy metal resistance |
| *rimP* | b3170 | 1.41305 | 0.486612 | Ribosome maturation factor involved in 30S subunit assembly |
| *rsxA* | b1627 | 1.643691 | 0.383259 | Controls oxidative stress response |
| *gloC* | b0927 | 1.401471 | 0.508469 | Involved in cellular stress response |

| | | | | |
|---|---|---|---|---|
| *hemF* | b2436 | 1.671551 | 0.535913 | Involved in heme biosynthesis |
| *ynfQ* | b4724 | 2.581637 | 0.751029 | Associated with anaerobic respiration |
| *rzpQ* | b1553 | 4.186201 | 1.32458 | Rac prophage protein; Unknown function |
| *alx* | b3088 | 2.187515 | 0.634505 | Involved in alkaline stress response |
| *ynfM* | b1596 | 1.802958 | 0.60589 | Associated with anaerobic respiration |
| *rrrQ* | b1554 | 2.701862 | 0.7954 | Rac prophage-encoded transcriptional regulator |
| *ydbJ* | b4529 | 3.362145 | 0.400845 | Predicted lipoprotein with unknown function |
| **C. Flagellar Assembly and Motility** | | | | |
| **Gene** | **Locus tag** | **log2(FC) P1 Vs C** | **log2(FC) O1 Vs C** | **Function** |
| *flgA* | b1072 | NS | 2.116683 | Flagellar assembly protein required for P-ring formation and motility |
| *flgB* | b1073 | NS | 2.694146 | Flagellar proximal rod protein required for basal body assembly and motility |
| *flgC* | b1074 | NS | 4.488955 | Flagellar distal rod protein required for |

| | | | | |
|---|---|---|---|---|
| | | | | basal body assembly and motility |
| *flgD* | b1075 | NS | 4.184013 | Flagellar hook capping protein required for hook assembly and motility |
| *flgE* | b1076 | NS | 3.173633 | Flagellar hook protein required for motility and flagellum assembly |
| *flgF* | b1077 | NS | 3.517814 | Flagellar proximal rod protein required for basal body assembly and motility |
| *flgG* | b1078 | NS | 2.93403 | Flagellar rod protein involved in basal body assembly and motility |
| *flgH* | b1079 | NS | 2.653642 | Flagellar L-ring protein required for basal body assembly and motility |
| *flgI* | b1080 | NS | 2.22738 | Flagellar P-ring protein required for basal body assembly and motility |
| *fliA* | b1922 | NS | 2.767232 | Alternative sigma factor regulating late flagellar gene expression and motility |

| | | | | |
|---|---|---|---|---|
| *fliF* | b1938 | NS | 3.137604 | Flagellar MS-ring protein required for flagellar assembly and motility |
| *fliG* | b1939 | NS | 2.820068 | Flagellar motor rotor protein; Involved in torque generation and motility |
| *fliH* | b1940 | NS | 3.174223 | Flagellar export protein involved in flagellar assembly |
| *fliM* | b1945 | NS | 4.409671 | Flagellar motor switch protein; Involved in motility and chemotaxis |
| *fliN* | b1946 | NS | 2.964226 | Flagellar motor switch protein involved in motility |
| **D. Anaerobic Respiration and Respiratory Metabolism** | | | | |
| **Gene** | **Locus tag** | **log2(FC) P1 Vs C** | **log2(FC) O1 Vs C** | **Function** |
| *frdD* | b4151 | 1.245561903 | 3.52604485 | Membrane anchor subunit of fumarate reductase; involved in anaerobic respiration |
| *fdnG* | b1474 | 2.742260763 | 5.2040039 | Involved in anaerobic respiration |
| *fdnH* | b1475 | 3.057166563 | 5.52917408 | Iron-sulfur electron transfer subunit involved in anaerobic respiration |

| | | | | |
|---|---|---|---|---|
| *fdnI* | b1476 | 1.918973966 | 3.33962753 | Involved in anaerobic respiration |
| *glpC* | b2243 | 1.07516691 | 2.46790928 | Respiratory electron transfer subunit involved in glycerol metabolism |
| *ynfE* | b1587 | 2.368333995 | 2.1457369 | Catalytic subunit of an anaerobic respiratory molybdoenzyme |
| *narY* | b1467 | 4.274622716 | NS | Involved in anaerobic respiration |
| *narZ* | b1468 | 3.456528656 | NS | Involved in anaerobic nitrate respiration |
| *narV* | b1465 | 3.541850603 | NS | Involved in anaerobic respiration |
| *ynfF* | b1588 | 2.053615418 | NS | Catalytic subunit of an anaerobic respiratory molybdoenzyme |
| *ynfG* | b1589 | 3.465860468 | NS | Electron transfer subunit of the YnfEFGH anaerobic reductase complex |
| *narG* | b1224 | NS | 6.74154569 | Involved in anaerobic nitrate respiration |
| *narH* | b1225 | NS | 6.6743613 | Iron-sulfur electron transfer subunit of nitrate reductase A |
| *narI* | b1227 | NS | 5.33486306 | Involved in electron transfer |
| *narJ* | b1226 | NS | 5.77108654 | Assembly chaperone required for nitrate |

| | | | | |
|---|---|---|---|---|
| | | | | reductase A maturation |
| *narK* | b1223 | NS | 6.23029614 | Nitrate/nitrite antiporter involved in anaerobic respiration |
| *napA* | b2206 | NS | 3.86619085 | Involved in anaerobic respiration |
| *napB* | b2203 | NS | 2.17746517 | Involved in anaerobic nitrate respiration |
| *napC* | b2202 | NS | 1.70380492 | Involved in anaerobic nitrate respiration |
| *frdA* | b4154 | NS | 2.69247556 | Involved in anaerobic respiration |
| *frdB* | b4153 | NS | 3.35919309 | Involved in anaerobic respiration |
| *frdC* | b4152 | NS | 3.36067487 | Involved in anaerobic respiration |
| *fdhF* | b4079 | NS | 1.81636338 | Involved in anaerobic formate metabolism |
| *dmsA* | b0894 | NS | 3.91993618 | Catalytic subunit of dimethyl sulfoxide reductase involved in anaerobic respiration |
| *dmsB* | b0895 | NS | 4.12028457 | Iron-sulfur electron transfer subunit of DMSO reductase involved in anaerobic respiration |
| *dmsC* | b0896 | NS | 3.25720119 | Membrane anchor subunit of DMSO reductase involved in anaerobic respiration |

| | | | | |
|---|---|---|---|---|
| *nirB* | b3365 | NS | 4.09036946 | Catalytic subunit of NADH-dependent nitrite reductase; Involved in anaerobic nitrite reduction |
| *nirD* | b3366 | NS | 4.88767751 | Small subunit of NADH-dependent nitrite reductase involved in anaerobic nitrite reduction |
| *nrfA* | b4070 | NS | 3.46265374 | Pentaheme cytochrome c nitrite reductase involved in anaerobic nitrite reduction |
| *nrfB* | b4071 | NS | 4.47541959 | Pentahaem c-type cytochrome electron transfer subunit of nitrite reductase system |
| *glpA* | b2241 | NS | 1.56602335 | Catalytic subunit of anaerobic glycerol-3-phosphate dehydrogenase involved in respiration |
| *glpB* | b2242 | NS | 2.04155368 | Membrane anchor subunit of anaerobic glycerol-3-phosphate dehydrogenase involved in respiration |

| | | | | |
|---|---|---|---|---|
| *hybA* | b2996 | NS | 1.92934348 | Iron-sulfur electron transfer subunit of hydrogenase-2 involved in anaerobic respiration |
| *hybB* | b2995 | NS | 2.23699979 | Membrane proton-translocating subunit of hydrogenase-2 involved in anaerobic respiration |
| *hybO* | b2997 | NS | 1.72641508 | Iron-sulfur electron transfer subunit of hydrogenase-2; Involved in anaerobic respiration |
| *dcuA* | b4138 | NS | 1.34322546 | Involved in anaerobic metabolism |
| *dcuB* | b4123 | NS | 2.66632463 | Fumarate/succinate antiporter involved in anaerobic respiration |
| *dcuC* | b0621 | NS | 3.63847582 | Anaerobic C4-dicarboxylate transporter; Involved in fumarate respiration |
| *fnr* | b1334 | NS | 10.8475992 | Global transcriptional regulator; Controls anaerobic metabolic response |
| *ccmA* | b2201 | NS | 1.88344823 | ATPase component of the cytochrome c |

| | | | | |
|---|---|---|---|---|
| | | | | maturation heme transporter |
| *ccmB* | b2200 | NS | 2.21136616 | Membrane component of the cytochrome c maturation system ; Involved in heme transport |
| *ccmC* | b2199 | NS | 1.62824943 | Component of the cytochrome c maturation system |
| *ccmF* | b2196 | NS | 1.33305475 | Membrane component of cytochrome c maturation system involved in heme attachment |
| **E. Stress response** | | | | |
| **Gene** | **Locus tag** | **log2(FC) P1 Vs C** | **log2(FC) O1 Vs C** | **Function** |
| *groS* | b4142 | - 0.972909266 | - 1.666860035 | Co-chaperonin assisting GroEL-mediated protein folding |
| *htpG* | b0473 | - 0.990865634 | - 1.271119493 | Involved in protein folding and stress response |
| *lexA* | b4043 | 1.790911436 | NS | Master transcriptional repressor of SOS DNA damage response |

| | | | | |
|---|---|---|---|---|
| *recA* | b2699 | 1.984961719 | NS | DNA recombinase; mediates homologous recombination and SOS response activation |
| *recN* | b2616 | 2.144106023 | NS | DNA repair protein involved in double-strand break repair |
| *dinB* | b0231 | 2.087156516 | NS | SOS response |
| *dinG* | b0799 | 1.696548244 | NS | ATP-dependent DNA helicase involved in DNA repair and genome stability |
| *uvrA* | b4058 | 2.784467096 | NS | DNA damage recognition; Nucleotide excision repair |
| *uvrB* | b0779 | 1.361349184 | NS | DNA damage recognition; Nucleotide excision repair |
| *uvrD* | b3813 | 1.403801388 | NS | Maintains genome stability |
| *ruvA* | b1861 | 1.515700377 | NS | DNA repair and homologous recombination |
| *ruvB* | b1860 | 2.207295992 | NS | Homologous recombination |
| *polB* | b0060 | 1.120034383 | NS | Involved in SOS-induced DNA repair |
| *umuC* | b1184 | 1.40387372 | NS | DNA damage response |

| | | | | |
|---|---|---|---|---|
| *puuR* | b1299 | 2.854683749 | NS | Transcriptional repressor of putrescine utilization operon |
| *puuA* | b1297 | 1.853693041 | NS | Initiates putrescine degradation |
| *puuB* | b1301 | 3.41224749 | NS | Involved in putrescine degradation |
| *puuC* | b1300 | 2.104524502 | NS | Involved in putrescine degradation |
| *puuD* | b1298 | 1.491466779 | NS | Involved in putrescine degradation |
| *puuE* | b1302 | 2.302485277 | NS | Involved in putrescine degradation |
| *puuP* | b1296 | 1.974136412 | NS | Putrescine transporter involved in putrescine uptake |
| *patD* | b1444 | 3.416594504 | NS | Aminotransferase involved in putrescine degradation |
| *sad* | b1525 | 1.942786674 | NS | Involved in putrescine and GABA metabolism |
| *paaX* | b1399 | 2.717791757 | NS | Transcriptional repressor of phenylacetic acid degradation operon |

| | | | | |
|---|---|---|---|---|
| *paaC* | b1390 | 2.877949022 | NS | Involved in phenylacetic acid degradation |
| *paaE* | b1392 | 2.478934479 | NS | Reductase subunit of the phenylacetyl-CoA epoxidase complex |
| *paaF* | b1393 | 3.39873352 | NS | Involved in phenylacetic acid degradation |
| *paaG* | b1394 | 3.344861588 | NS | Involved in phenylacetic acid degradation |
| *paaH* | b1395 | 2.057845824 | NS | Involved in phenylacetic acid degradation |
| *paaK* | b1398 | 2.953274647 | NS | Initiates phenylacetic acid degradation |
| *paaZ* | b1387 | 1.749811973 | NS | Involved in phenylacetic acid degradation |
| *hdeB* | b3509 | -6.032926125 | NS | Acid stress response |
| *mgrB* | b1826 | -1.58958975 | NS | Membrane-bound negative regulator of PHoP/PhoQ two-component system |
| *focA* | b0904 | -1.446705416 | NS | Bidirectional formate channel involved in anaerobic metabolism |
| *hchA* | b1967 | -1.467717915 | -1.612039167 | Stress response |

| | | | | |
|---|---|---|---|---|
| *appY* | b0564 | -4.270956711 | NS | Anaerobic stress regulation |
| *evgA* | b2369 | -1.583163986 | NS | Controls acid resistance and multidrug resistance |
| *gadX* | b3516 | -2.697256469 | NS | Regulates glutamate-dependent acid resistance |
| *fis* | b3261 | -2.024034581 | NS | Global transcriptional regulation; Controls chromosome organization and gene expression |
| *sodB* | b1656 | 1.420746313 | NS | Oxidative stress response |
| *azoR* | b1412 | 3.417350449 | NS | Protection against oxidative stress |
| *bssR* | b0836 | -1.882062703 | 5.488143434 | Acts as iron and zinc sensor; Regulates genes responsible for metal homeostasis, acid tolerance and biofilm formation |
| *dnaK* | b0014 | NS | -1.222313482 | Heat shock protein; prevents protein aggregation |
| *groL* | b4143 | NS | -1.295839196 | Mediates protein folding |
| *ibpA* | b3687 | NS | -2.455204723 | Small heat-shock protein; Act as ATP-independent molecular chaperone |

| | | | | |
|---|---|---|---|---|
| *ibpB* | b3686 | NS | -3.337624137 | Small heat-shock protein; Protects against severe environmental stress |
| *hslU* | b3931 | NS | -1.257455799 | ATP-dependent unfoldase of HslVU protease complex |
| *hslV* | b3932 | NS | -1.262495638 | Protein degradation |
| *cspA* | b3556 | NS | 1.920145326 | Cold-shock response, Promotes translation and transcription antitermination |

***F.* Significantly enriched regulators involved in core response to *racR* silencing**

| Regulator | log2(FC) P1 Vs C | log2(FC) O1 Vs C | Function |
|---|---|---|---|
| RacR | 10.299 | 15.312 | Rac prophage repressor; Suppresses toxic ydaS-ydaT operon |
| PspF | 7.724 | NS | Transcriptional activator of Phage shock response |
| LexA | 3.727 | NS | Repressor of SOS DNA-damage response |
| FliZ | -4.883 | NS | Promotes flagellar motility and repressing stress response |
| GadW | -7.813 | NS | Controls glutamate-dependent acid |

| | | | |
|---|---|---|---|
| | | | resistance (GADR) system |
| AdiY | -6.104 | NS | Transcriptional activator of arginine-dependent acid resistance system |
| TorR | -5.58 | NS | Controls TMAO-dependent anaerobic respiration |
| GadR | -5.425 | NS | Acid resistance |
| YdeO | -4.883 | NS | Controls genes involved in acid resistance, anaerobic respiration, SOS response |
| GadE | -3.734 | NS | Glutamate-dependent acid resistance |
| GadX | -3.734 | NS | Glutamate-dependent acid resistance |
| PhoP | -2.279 | NS | Controls gene expression essential for bacterial survival and virulence |
| QseB | -4.883 | NS | Controls motility, virulence, antibiotic tolerance |
| H-NS | -2.247 | -2.076 | Global transcriptional repressor |
| BluR | -9.766 | -18.375 | Biofilm maturation and regulates Rcs stress response |

| | | | |
|---|---|---|---|
| McbR | -9.766 | -9.188 | Biofilm formation, colanic acid production, and bacterial virulence |
| CsrA | -3.662 | NS | Global post-transcriptional regulator; controls carbon metabolism, motility, biofilm formation |
| CysB | -4.439 | NS | LysR-family transcriptional regulator of sulfate assimilation and cysteine biosynthesis |
| GlaR | -2.035 | NS | Transcriptional regulator of carbon starvation and lysine degradation |
| ppGpp | -1.585 | NS | Global transcriptional regulator |
| DcuR | NS | 8.481 | Controls C4-dicarboxylate transport and anaerobic fumarate respiration |
| NarP | NS | 8.287 | Controls anaerobic respiration |
| RstA | NS | 7.35 | Regulates acid tolerance, biofilm formation, stress response |

| | | | |
|---|---|---|---|
| NarL | NS | 7.146 | Response regulator controlling nitrate and nitrite respiration gene expression |
| ModE | NS | 5.992 | Governs molybdenum metabolism |
| IscR | NS | 3.898 | Maintains Iron–sulfur homeostasis |
| FNR | NS | 3.765 | Global oxygen-responsive transcriptional regulator; controls aerobic and anaerobic metabolism |
| NikR | NS | 15.312 | Maintains intracellular nickel homeostasis |
| FlhDC | NS | 7.113 | Flagellar biosynthesis, governs virulence and swarming motility |
| IHF | NS | 1.869 | Nucleoid-associated DNA-binding protein |
| StpA | NS | -7.875 | Global gene silencing, analogue of H-N-S |

List of RNA-seq–derived log2 fold changes (log2FC) for selected genes in the P1 Vs control (C) and O1 Vs control (C) comparisons after CRISPRi-mediated silencing of *racR*. Positive log2FC values indicate upregulation and negative values indicate downregulation relative to the control strain. Sections are organized by functional group: (A) Coordinated expression of Rac prophage genes upon *racR* silencing; (B) Genes commonly upregulated in both P1 Vs C and O1 Vs C comparisons (*racR* genes excluded) (C) Flagellar assembly and motility (D) Anaerobic respiration and respiratory metabolism; (E) Stress response (F) Significantly enriched regulators implicated in the core response to *racR* silencing. For each gene, the table provides gene name/locus, log2FC (P1 Vs C), log2FC (O1 Vs C), and annotated functions (Sources: Supplementary data I and II). This dataset was used to generate the heat map across different categories of processes.
